# The sequence context of RG/RGG motifs determines condensate formation, transportin-1 binding and chaperoning

**DOI:** 10.64898/2026.01.26.701751

**Authors:** Sinem Usluer, Yukti Khanna, Saskia Hutten, Đesika Kolarić, Benjamin Bourgeois, Iva Pritišanac, Dorothee Dormann, Tobias Madl

## Abstract

Intrinsically disordered arginine-glycine-rich (RG/RGG) regions are highly abundant in the eukaryotic proteome. Proteins containing these motifs participate in fundamental cellular processes, including nuclear import, transcriptional regulation, biomolecular condensate formation, and apoptosis. Mutations or dysfunction of RG/RGG proteins have been implicated in neurodegenerative diseases and cancer. Although some RG/RGG proteins have been shown to drive condensate formation, localize to membrane-less organelles, interact with nuclear import receptors, or undergo arginine methylation, these properties are not shared uniformly across the proteome. The considerable diversity in RG/RGG motif length and amino acid composition raises the question of which sequence features determine their functional behaviour. To address this, we conducted a systematic bioinformatics and experimental analysis, combining synthetic and natural peptides with studies on the RNA-binding protein FUS as a model system. Our results reveal that the sequence composition of RG/RGG motifs is a key determinant of their capacity for RNA-mediated condensate formation, stress granule recruitment, and transportin-1–mediated chaperoning and nuclear import. These findings provide new insight into the sequence grammar of disordered RG/RGG regions and how it encodes the multifunctionality of these proteins in cellular regulation.

## INTRODUCTION

Eukaryotic cells organize their intracellular components into membrane-bound organelles as well as membrane-less organelles (MLOs). These MLOs can be found in the nucleus, for example the nucleolus, nuclear speckles, promyelocytic leukemia protein nuclear bodies (PML-NBs), and Cajal bodies, as well as in the cytosol, for example, stress granules (SGs) and P-bodies (1,2). These compartments provide specific environments, for biochemical reactions (3,4), or spatial and temporal storage of specific biomolecules in response to various stress conditions (3). MLOs are involved in the regulation of a variety of cellular processes like gene expression, stress response, signal transduction, and development (3,5,6) Dysregulation of MLO assembly is associated with a large number of diseases, including neurodegenerative diseases and cancers (5,7,8).

MLO formation is regulated by condensation driven by weak and multivalent protein-protein and protein-nucleic acid interactions involving intrinsically disordered regions (IDRs), DNA/RNA-binding domains, and oligomerization domains (3,9–12). IDRs are highly abundant in the human proteome and do not adopt any stable secondary or tertiary structure (13–17). This allows them to act as hubs in signaling networks and to mediate protein-protein interactions in a plethora of biological processes which include, for example, signaling, replication, cell-cycle control, transcription, and chaperoning (13–17). IDRs frequently harbor low-complexity regions, which are highly enriched in certain amino acids, such as FG repeats, polyglutamine, polyglutamine/asparagine-rich repeats, or arginine-glycine/-glycine (RG/RGG) repeats (18). Importantly, RG/RGG regions are not defined by the RG dipeptide alone: proteome-wide context mapping has shown that RG motifs occur in distinct local sequence environments and can be distinguished by contextual/compositional signatures, and machine-learning “grammar” approaches similarly identify short sequence features (e.g., enriched 2–3 mer patterns) that help predict RNA-granule–associated proteins, collectively arguing that additional residues and local motif context contribute to RG/RGG-linked condensate biology (19,20). Prion-like domains (PrLDs) are a subset of low-complexity IDRs and are frequently linked to aberrant self-association. Several studies have recently underlined the essential role of disordered RG/RGG regions in the regulation of condensate and MLO formation by promoting multivalent homo- and/or heterotypic protein-protein and protein-RNA interactions (3,9,21). The RNA-binding behaviour of core RGG/RG motifs can be reshaped by neighbouring sequence elements, indicating that residues flanking RGG/RG motifs can tune affinity and/or selectivity rather than acting as interchangeable repeats (22). RG/RGG regions are enriched in RBPs, such as Fused-in-sarcoma (FUS), Ewing sarcoma breakpoint region 1 protein (EWSR1), TATA-binding protein-associated factor 2N (TAF15), Heterogeneous nuclear ribonucleoprotein A1 (hnRNPA1), Cytotoxic granule associated RNA binding protein TIA1 and Src-associated in mitosis 68 kDa protein (1,23). Proteins with RG/RGG regions are involved in various biological processes, including translation regulation, apoptosis, and DNA damage signaling; and they have been implicated in several human diseases, including neurodegeneration, cancer and infectious diseases (8). For example, mutations in the disordered prion-like domains of hnRNPA1 have been described to cause self-aggregation, fibrillation, and association with SGs in amyotrophic lateral sclerosis (ALS) (24). Also, FUS hypomethylated in RG/RGG regions is enriched in cytosolic inclusions in frontotemporal dementia (FTD) patients (25,26). Recent evidence further highlights that arginine methylation within FUS RGG domains can qualitatively rewire phase behaviour by shifting the balance between homotypic versus heterotypic interactions, reinforcing that the regulatory ‘context’ of RG/RGG regions includes both neighbouring residues and their modification state (27).

Beyond condensate formation, RG/RGG regions can contribute to protein nuclear import by serving as nuclear localization signal (NLS) and binding to nuclear import receptors (NIRs) (25,28). Moreover, via binding to RG/RGG regions, NIRs can act as chaperones, preventing the formation of condensates and pathological aggregates (28–33). This chaperone-like activity of nuclear import receptors has recently been synthesized into a broader ‘gatekeeper’ framework for ALS/FTD-linked RBPs, emphasizing that NIRs can both promote nuclear localization and antagonize pathological phase transitions of prion-like RBPs (34). In FUS, TNPO1 recognizes a C-terminal PYNLS, and ALS-linked mutations in this PY-NLS impair TNPO1-dependent import, promoting cytoplasmic accumulation and reduced chaperoning (7,31,35). The RG/RGG motifs of FUS lie within the N-terminal low-complexity region upstream of the PY-NLS, where they contribute to condensation and can be engaged by nuclear import receptors during chaperoning (28,32,35). Beyond TNPO1, other NIRs, including transportin-3, importin β, importin 7 and importin β/7, have been shown to suppress phase separation and SG association of FUS by binding to its RG/RGG regions (33). Moreover, TNPO1 can suppress phase separation of other RBPs containing RG/RGG regions, including FUS, TAF15, EWSR1, hnRNPA1, hnRNPA2 and Cold-inducible RNA-binding protein (CIRBP) (28,33,36).

In light of those examples, it is becoming evident that RG/RGG regions are key regulatory elements for condensate and MLO formation, as well as for binding and mediating import and chaperoning by NIRs (3,21). Despite these advances, we still lack mechanistic rules that connect specific local sequence features within RG/RGG regions, especially residues immediately adjacent to RG motifs and their modification state, to RNA-driven condensation versus aggregation and to TNPO1 binding (20,27,34). Contrary to the general assumption that all RG/RGG proteins phase separate, some RG/RGG proteins, such as Histone-lysine N-methyltransferase 2B, do not phase separate nor localize to any MLOs in cells (PhaSepDB2.0). Moreover, artificial mutations in the RG/RGG region of LAF1 and FUS alter their phase separation behavior (30,37), with arginine to lysine mutations decreasing or abolishing phase separation, respectively (30,37). Here, we map how residues surrounding and immediately preceding RG/RGG motifs are distributed across human disordered RG/RGG regions and relate this local context to tract organization and functional outcomes. It therefore seems possible that condensate formation, TNPO1 binding, chaperoning and nuclear import of RG/RGG proteins could be encoded in the sequence context of RG/RGG proteins. Thus, we hypothesized that the nature of the preceding residue “[X]” of [X]RG/[X]RGG repeats fine-tunes the phase separation behavior and association with TNPO1 and therefore investigated how the sequence context of RG/RGG regions determines phase separation in vitro, association with condensates in cells, TNPO1-binding, chaperoning, and nuclear import.

In this study, we dissect how local sequence context around RG/RGG motifs tunes RNA-driven condensation and recognition by the nuclear import receptor TNPO1. First, we establish a minimal and biologically relevant framework for studying RG/RGG repeats: combining coarse-grained simulations with in vitro assays, we show that RNA strongly amplifies residue-specific differences at the position immediately preceding RG, and we use this rationale to focus on short [XRGG] repeat peptides. Second, we place these model systems in a proteome-scale context by mapping how RG/RGG motifs are organized in human disordered regions and find that RG motifs in canonical RG/RGG proteins are typically concentrated into compact tracts separated by short linkers. Building on this foundation, we systematically test how charge and aromatic content in model peptides and natural RG/RGG segments shape condensate versus aggregate formation, and how these same sequence features modulate TNPO1 binding and chaperoning and influence stress granule recruitment and nuclear import in cells.

## MATERIAL AND METHODS

### Plasmids and Synthetic Peptides

We generated and cloned optimized cDNA expression constructs (Genscript) into the His_6_-protein A pETM11 vector, which harbors a tobacco etch virus (TEV) protease cleavage site after the protein A tag, using NcoI/BamHI restriction digest. These constructs were generated for fragments of human TNPO1 (Q92973; residues 1 to 898), human FUS (P35637; residues 454 to 526; named FUS^RGG3-PY^), and human FUS variants harboring mutations at positions 472, 480, 486, 490, 494, and 502, to E, F, R, Y, T and C, respectively. GCR_2_-GFP_2_-FUS wild type and mutant human expression constructs were generated by cloning optimized cDNAs into a pEGFP-C1 vector containing a GCR_2_-GFP_2_-cassette, using EcoRV and BamHI.

Synthetic peptides, which are listed in a supplementary excel sheet, were synthesized by Peptide Specialty Laboratories GmbH, Heidelberg, Germany. The peptides were HPLC-purified to a final purity above 95 %. They were dissolved in a buffer containing 50 mM Tris-HCl, 150 mM NaCl, 0.04 % (w/v) NaN_3,_ pH 7.5 and with or without 2 mM TCEP (2-carboxyethyl)phosphine), depending on the experiment. For SG recruitment assays, peptide concentration was adjusted by measuring fluorescence intensity at 480 nm using a Nanodrop device (Thermo).

### Protein Expression and Purification

For expression of recombinant, unlabeled His_6_-protein A-TNPO1 (pETM11-TNPO1), the respective bacterial expression vector was transformed into Escherichia coli (E.coli) BL21-DE3 Star strain, one colony was picked for each construct and grown in 20 mL lysogeny broth (LB) medium for 16 hours at 37°C. Next, 1 mL of pre-culture was grown for 3 days in 1L minimal medium containing 100 mM KH_2_PO_4_, 50 mM K_2_HPO_4_, 60 mM Na_2_HPO_4_, 14 mM K_2_SO_4_, 5 mM MgCl_2_and pH 7.2 adjusted with HCl and NaOH with 0.1 dilution of trace element solution (41 mM CaCl_2_, 22 mM FeSO_4_, 6 mM MnCl_2_, 3 mM CoCl_2_, 1 mM ZnSO_4_, 0.1 mM CuCl_2_, 0.2 mM (NH_4_)_6_Mo_7_O_24_, 17 mM EDTA, supplemented with 6 g of glucose (Carl Roth, Karlsruhe, Germany) and 3 g of NH_4_Cl (Carl Roth, Karlsruhe, Germany) at 30°C. Cells were diluted to an OD (600nm) of 0.8 and induced with 0.5 mM Isopropyl β-D-1-thiogalactopyranoside (IPTG, BLDpharm, Shanghai, China), followed by protein expression for 6 h at 30°C. For the expression of recombinant, unlabeled His_6_-protein A-FUS^RGG3-PY^ and FUS^RGG3-PY^ mutants, the respective bacterial expression vector was transformed into the E.coli BL21DE3 Star strain, which was plated onto a LB agar plate. Pre-culture was started from a single isolated colony from plate grown in 20 mL LB medium for 16 hours at 37°C. Next, 10 mL of pre-culture was grown in 1L LB medium, and cells were induced at an OD (600 nm) of 0.8 with 0.5 mM IPTG, followed by protein expression for 16 hours at 20°C. Cell pellets corresponding to protein expression of unlabeled disordered protein fragments (FUS^RGG3-PY^ and FUS^RGG3-PY^ mutants) were harvested and sonicated in denaturating lysis buffer (50 mM Tris-HCl, 150 mM NaCl, 20 mM imidazole, 6 M urea, pH 7.5), whereas folded proteins fragments (TNPO1) were harvested and sonicated in non-denaturating lysis buffer (50 mM Tris-HCl, 150 mM NaCl, 20 mM imidazole, 2 mM TCEP, pH 7.5). His_6_-protein A-tagged recombinant proteins were then purified using Ni-NTA agarose beads (Qiagen, Hilden, Germany) in elution buffer (50 mM Tris, 1000 mM NaCl, 500 mM imidazole, 2 mM TCEP, pH 7.5). Eluted His_6_-protein A-tagged recombinant proteins were subjected to 2% (w/w) His_6_-tagged TEV protease cleavage overnight at 4°C. TEV-cleaved recombinant proteins were separated from the His_6_-protein A tag using a second step of Ni-NTA purification. Cleaved recombinant proteins were desalted into low imidazole buffer (50 mM Tris-HCI, 150 mM NaCl, 20 mM imidazole, 2 mM TCEP, pH 7.5) using a desalting column (HiPrep 26/10, GE Healthcare, Chicago, IL, USA) on an ÄKTA Pure system (GE Healthcare, Chicago, IL, USA). A final size exclusion chromatography purification step was performed in the buffer of interest (50 mM Tris·HCl, 150 mM NaCl, 2 mM TCEP, 0.04 % (w/v) NaN_3_, pH 7.5). The TNPO1 sample was purified using a Hiload 16/600 Superdex 200 pg gel filtration column, GE Healthcare (Chicago, IL, USA). FUS^RGG3-PY^, X6C FUS^RGG3-PY^, X6E FUS^RGG3-PY^and X6T FUS^RGG3-PY^ were purified using a Superdex 75 gel filtration column GE Healthcare (Chicago, IL, USA). After elution of His_6_-protein AX6R FUS^RGG3-PY^, X6F FUS^RGG3-PY^, and X6Y FUS^RGG3-PY^ recombinant proteins from nickel columns, DNase and RNase treatments were performed for 30 minutes at room temperature. The proteins were then cleaved by the addition of 2 % (w/w) His_6_-tagged TEV for 1 hour at room temperature. The proteins were uploaded to a HiTrap Heparin column GE Healthcare (Chicago, IL, USA) and eluted with 50 mM Tris-HCl, 1000 mM NaCl, 6 M urea, 20 mM imidazole, 2 mM TCEP, pH 7.5. On the same day, the proteins were desalted into 50 mM Tris·HCl,150 mM NaCl, 2 mM TCEP, 0.04 % (w/v) NaN_3_, pH 7.5 using a HiTrap Desalting column GE Healthcare (Chicago, IL, USA). Protein concentrations were estimated based on their absorbance at 280 nm, assuming that the ε value at 280 nm was equal to the theoretical ε value.

### Isothermal Titration Calorimetry (ITC)

All protein samples and peptides were equilibrated in the same buffer containing 50 mM Tris-HCl, 150 mM NaCl, 2 mM TCEP, 0.04 % (w/v) NaN_3_, pH 7.5. ITC measurements were carried out on a VP-ITC instrument (Malvern, United Kingdom) with 28 rounds of 10-μL injections at 25 °C. The time between injections is 210 seconds. However, the time between injections was increased to 240 seconds for [LRGG]_7_, FUS^RGG3-PY^ and X6R_ FUS^RGG3-PY^ to allow the system to reach equilibration. Integration of peaks corresponding to each injection, subtraction of the contribution of protein dilution, and correction for the baseline were performed using the Origin-based 7.0 software provided by the manufacturer. The equilibrium binding constant (Ka) and enthalpy of the complex formation (ΔH) were determined by curve fitting using a standard one-site model, except for [RRGG]_7_, for which a two-binding site model was used.

### Turbidity Assays

All [XRGG]_7_ and RNA (12 × UG repeats, Eurofins, purification by desalting) samples were prepared in the buffer containing 50 mM Tris-HCl, 150 mM NaCl, 0.04 % (w/v) NaN_3_, pH 7.5, while FUS^RGG3-PY^/FUS^RGG3-PY^ mutants, natural RGG peptides, and RNA (12 × UG repeats) samples were prepared in the buffer containing 50 mM Tris-HCl, 150 mM NaCl, 2 mM TCEP, 0.04 % (w/v) NaN_3_, pH 7.5. Turbidity measurements were conducted at 620 nm in 96-well plates with 90-μL samples using a BioTek Power Wave HT plate reader (BioTek, Santa Clara, California, USA). Measurements were performed directly after pipetting.

### Differential Interference Contrast Microscopy

All [XRGG]_7_/RNA (12 × UG repeats) samples were prepared in the buffer containing 50 mM Tris-HCl, 150 mM NaCl, 0.04 % NaN_3_ pH 7.5, while FUS^RGG3-PY^, FUS^RGG3-PY^ mutants, natural RGG peptides, and RNA (12 × UG repeats) samples were prepared in a buffer containing 50 mM Tris-HCl, 150 mM NaCl, 2 mM TCEP, 0.04 % NaN_3_, pH 7.5. Droplet formation was induced by the addition of a 1:2 or 1:3 ratio for RNA:peptide/protein depending on the phase separation propensity of peptide/protein. The 25 μL samples were plated on a 30-mm No. 1 round glass coverslip and mounted on an Observer D1 microscope with a 100×/1.45 oil immersion objective (Zeiss, Oberkochen, Germany). Protein droplets were viewed using a HAL 100 halogen lamp, and images were captured with an OrcaD2 camera (Hamamatsu, Shizuoka, Japan) using VisiView 4.0.0.13 software (Visitron Systems GmbH, Puchheim, Germany). Images were recorded 30 minutes after the addition of RNA. Images were recorded without the addition of RNA for aggregation-prone peptides. The microscope images were processed using Fiji/ImageJ 1.53a software (Bethesda, US), applying a linear enhancement for brightness and contrast.

### Fluorescence Polarization

N-terminally fluorescein isothiocyanate (FITC)-labeled FUS^PY^ was dissolved in a buffer containing 50 mM Tris, 150 mM NaCI, 2 mM TCEP, and 0.04 % (w/v) NaN_3_ pH 7.5. Measurements were performed in black 384-well plates at room temperature using a ClarioStar Plus (BMG labtech, Ortenberg, Germany) spectrophotometer. Filters were selected as a function of FITC optical characteristics (λex= 495 nm, and λem= 530 nm). 100 nM FITC-labelled FUS^PY^ was incubated in either increasing concentration of TNPO1 or constant concentration (90 µM) of [RRGG]_7_ with increasing concentration of TNPO1, in a final volume of 35 µl. The polarization data were fitted using GraphPad prism 8 with the following equation:

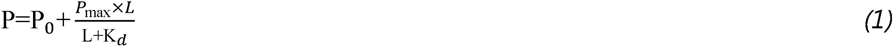

Here, P_0_ represents the polarization of FITC-labelled peptides in the absence of TNPO1, and P_max_ to the highest polarization of the binding curve corresponding to the saturation of the interaction. L corresponds to the concentration of TNPO1 protein, and K_d_ is the dissociation constant.

### SG association assay in semi-permeabilized cells

HeLa cells were grown on poly-L-lysine–coated 12-mm coverslips No. 1.5. In brief, SGs were induced by 3h treatment with 10 µM MG132 before the plasma membrane was selectively permeabilized using 0.003 to 0.005% digitonin in potassium phosphate buffer (KPB) (20 mM potassium phosphate, pH 7.4, 5 mM MgCl2, 200 mM KOAc, 1 mM ethylene glycol tetraacetic acid, 2 mM dithiothreitol (DTT), 1 μg/mL each aprotinin, pepstatin, and leupeptin) and all soluble cytoplasmic components were washed out. After transport passages through the nuclear pores were blocked by WGA, cells were incubated for 30 min with 1 µM peptide in KPB. After several stringent washes, cells were fixed and SGs were visualized by immunostaining of G3BP1. SG recruitment of RGG peptides was analyzed with an inverted Leica SP8 microscope and the LAS X imaging program (Leica), using lasers for 405 nm, 488 nm and 552 nm excitation. Images were acquired using two-fold frame averaging with a 63x 1.4 oil objective and an image pixel size of 59 nm. The following fluorescence settings were used for detection: DAPI: 419-442 nm, GFP: 498-533 nm, Alexa 555: 562-598 nm. Recording was performed sequentially to avoid bleed-through using a conventional photomultiplier tube. Images were processed and analyzed using Image J/Fiji software, applying linear enhancement for brightness and contrast and implemented plugins for measurement of pixel intensities in SGs. Statistical analyses were performed in GraphPad Prism 9.

### Hormone induced nuclear import assay

HeLa cells were grown for at least 2 passages in DMEM supplemented with 10% dialyzed FBS. Cells were transfected with GCR_2_GFP_2_-FUS reporters using Lipofectamine 2000. 20h later nuclear import of the constructs was induced by the addition of dexamethasone (5 µM final concentration) in imaging medium (Fluorobrite; Gibco) supplemented with 10% dialyzed FBS. Nuclei were stained using Draq5 (Thermo) for 30 min before images were acquired for a duration of 30 min in 2.5 min intervals in an environmental chamber at 37°C and 5% CO_2_ using an inverted spinning disc microscope (5 Elements, Visitron) equipped with a confocal spinning disc (CSU-W1; Yokogawa, Tokyo, Japan) and a 60x/1.2 water immersion lens. Images were acquired using the 488 nm and 650 nm laser line and 2 sCMOS cameras for simultaneous acquisitons at bin 1×1. Analysis was performed using Fiji. If applicable, the StackReg Plugin was used to compensate xy drift of cells over time before downstream analysis. To determine nuclear import nuclear, at least 19 cells per replicate was analyzed by measuring the fluorescence intensity of the nuclei (as stained by Draq5) over a representative circular region of interest in the cytoplasm (C). Background subtracted values were then plotted as N/C ratio over time.

### Image Processing

The microscopy images were processed using Fiji/ImageJ 1.53a software (Bethesda, US), applying linear enhancement for brightness and contrast.

### Bioinformatics Studies

To gain additional insight into the occurrence of RG/RGG regions across the human IDRome, we quantified them using an in-house Python script (https://github.com/DesikaKolaric/RG-RGG-proteins). First, we predicted intrinsically disordered regions (IDRs) from all canonical amino acid sequences in the human proteome (downloaded from UniProt, ProteomeID UP000005640, August 2019) using the state-of-the-art sequence-based predictor of intrinsic disorder, SPOT-Disorder (38). We obtained 21,252 human intrinsically disordered regions, defined as disordered regions of 30 or more residues. Subsequently, we identified all RG/RGG repeats with a spacing of 0-5 residues in each IDR sequence using a regular expression of the form RG[G].(0,5)RG[G] (Python standard module RegEx). To explore the distribution of consecutive RG/RGG repeats, a stepwise search was conducted, beginning with two repeats and incrementally increasing the count up to twelve. To define the RG/RGG repeat region, all residues between the first arginine of the motif and the last glycine of the last motif were considered, as well as ten residues preceding the first arginine and ten residues following the last occurrence of the RG/RGG repeat region. Next, we considered the frequency of each amino acid in regions of 3, 4, and 5 consecutive RG/RGG repeats, by counting the occurrences of each amino acid within the repeat region and dividing it by the total number of all other residues in the region. The same protocol was used to investigate the frequency of each amino acid in RG/RGG regions in IDRs of proteins that are known to localize in stress granules (39) as one set and of proteins known to bind to TNPO1 (40,41) as another set.

### Coarse-Grained Slab Simulations of [XRGG]₇ Peptides With and Without RNA

To quantify RNA-dependent phase separation of [XRGG]₇ peptides, we performed coarse-grained slab simulations using the CALVADOS 3 model (42–44). Simulation inputs were generated via custom Python scripts based on the CALVADOS configuration framework (prepare_slab_idr.py for peptide-only systems and prepare_slab_mix.py for peptide+RNA systems). Peptide-only simulations contained 200 copies of a single [XRGG]₇ variant, whereas peptide–RNA simulations included 200 peptide chains and 60 molecules of a non-specific [UG]₁₂ RNA.

Protein termini were modeled as charged at both N- and C-termini (charge_termini=’both’). All systems were simulated in a slab geometry (topol=’slab’) within a periodic simulation box of 15 × 15 × 80 nm. Simulations were conducted at 298.15 K, pH 7.0, and 150 mM ionic strength, using GPU acceleration (platform=’CUDA’).

Trajectory snapshots were recorded every 100,000 integration steps (wfreq = 100000), with 1,000 frames saved per replica (N_frames = 1000), corresponding to one snapshot per nanosecond. Each simulation included a slab equilibration phase of 10 million steps (steps_eq = 10,000,000), followed by a 100 million-step production run (steps = 100,000,000). Three independent replicas were performed for each condition.

Peptide and RNA sequences were provided in a FASTA file (input/xrgg.fasta), and residue parameters were defined in input/residues_C2RNA.csv. RNA parameters used in the Components configuration were: rna_kb1 = 1400.0, rna_kb2 = 2200.0, rna_ka = 4.20, rna_pa = 3.14; excluded volume settings were rna_nb_sigma = 0.4, rna_nb_scale = 15, and rna_nb_cutoff = 2.0. The preparation and adjacent files are available on https://github.com/yukti-khanna/rg-rich-regions.

### Trajectory Analysis and Quantification of Partition Coefficients

Simulation trajectories were analyzed using the python scripts modified from CALVADOS SlabAnalysis module embedded in the run configuration (42,44). Trajectories were centered from frame 250 onward to minimize early transient behavior (SlabAnalysis.center(start=250, center_target=’all’)). One-dimensional concentration profiles along the slab axis were computed (SlabAnalysis.calc_profiles()), and dense- and dilute-phase concentrations were estimated using SlabAnalysis.calc_concentrations(pden=1.0, pdil=5.0), which defines the regions based on default interface handling with the specified multipliers.

Outputs included per-component concentration profiles (*_profile.npy) and a summary file (*_ps_results.csv) containing estimated concentrations in the dense and dilute phases (reported in mM), along with corresponding integration boundaries (cutoffs_dense_left/right, cutoffs_dilute_left/right).

Partition coefficients were calculated as:

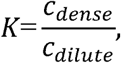

separately for peptides and RNA. Values were extracted from *_ps_results.csv using a custom script (scan_partition_coefficients.py), which excluded simulations where concentrations were non-finite or ≤0. For each condition, the mean, standard deviation (SD; with ddof=1), and standard error of the mean (SEM) were computed across replicas for both *K* and *log*_10_ *K*. Profile plots, partitioning statistics, and time series of phase concentrations were generated using scripts available at https://github.com/yukti-khanna/rg-rich-regions.

### RG motif counts and inter-RG linker lengths

RG motifs were defined as the dipeptide “RG” (case-normalized by converting sequences to uppercase). For each protein sequence in the predicted IDR proteome FASTA and the curated RG/RGG protein FASTA (Supplementary File 1, 2), we counted RG motifs as seq.count(“RG”) and compiled the distribution of RG motifs per protein. For plotting (Fig. 3A), counts were clipped to a maximum bin (30) and converted to percent of proteins per bin.

To quantify spacing between consecutive RG motifs, we identified all RG motif start positions using regular-expression matching (re.finditer(“RG”, seq)) and computed the inter-RG linker length as the number of residues between successive RG motifs:

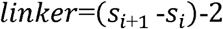

where and _+_ are consecutive RG start indices and “2” accounts for the RG dipeptide length. Linkers 0 were pooled across all sequences in each dataset and plotted as histograms over 0–26 residues (Fig. 3B). Scripts for plotting is available at https://github.com/yukti-khanna/rg-rich-regions.

### Sliding-window RG organisation, KDE contours, and compact tract calling

To quantify local RG organisation, each sequence was scanned with sliding windows of 20, 40, 60, 80, and 100 residues. Windows containing at least two RG motifs were retained, and for each window we recorded: (i) the number of RG motifs per window and (ii) the mean inter-RG linker length within that window (computed from consecutive RG start positions using the linker definition above). These paired measurements formed the inputs for the 2D analyses in Fig. 3C–D (with contour boundaries for all window sizes reported in Supplementary File 5).

For each dataset (known RG/RGG proteins (7) or all predicted IDRs - Supplementary file 1 and 2) and window size, we estimated the joint density of (mean linker length, RG motifs per window) using a 2D Gaussian KDE (scipy.stats.gaussian_kde). The KDE was evaluated on a 100×100 grid spanning the observed x/y ranges. For visualization, densities were additionally scaled by the maximum grid density to yield a 0–1 “scaled density” heatmap. To draw cumulative contours, grid densities were flattened and sorted; the normalized cumulative sum over sorted density values was used to select density thresholds corresponding to 0.7, 0.8, and 0.9 cumulative levels, which were then plotted as the 70/80/90% contour lines (Fig. 3C–D). Vertical guide lines at 7 and 15 residues were overlaid to indicate the compact and permissive spacing thresholds.

Guided by these distributions, compact RG tracts were defined by grouping consecutive RG motifs along each sequence using an operational two-threshold rule: motifs were assigned to the same tract if separated by ≤7 residues, with a more permissive ≤15-residue gap allowed only to bridge otherwise compact segments (i.e., to avoid splitting a tract due to a single moderately longer spacer between short-linker clusters). Tract sequences and identifiers were exported for downstream summaries and plotting (e.g., tract length vs RG count). Scripts for calculation and plotting is available at https://github.com/yukti-khanna/rg-rich-regions.

## RESULTS

### X residues determine condensate formation of XRGG peptides

Abnormal phase separation or mutations in RBPs containing RG/RGG regions leads to the aggregate formation in the cytoplasm and is linked to neurodegenerative diseases (45–49). The residues surrounding and preceding the arginines of RG/RGG regions vary, yet their role in regulating phase separation is unclear, as well as how this local sequence context tunes RNA-driven condensation. We therefore asked whether the residue immediately preceding the RG motif (the “X” position in [X]RG/[X]RGG) encodes systematic differences in RNA-driven assembly. We selected [XRGG]_7_ as a minimal model because short RG/RGG repeat arrays predominate in human IDRs (shown below). To directly test how X affects condensation, we performed coarse-grained slab simulations (CALVADOS 3) for a library of [XRGG]_7_ peptides in the presence and absence of non-specific [UG]_12_ RNA (Figure 1) (42,43) (Supplementary File 3). In peptide-only simulations, [XRGG]_7_ variants behaved similarly and did not form a robust condensed phase under these conditions (Figure 1A). In contrast, addition of RNA uncovered strong sequence dependence: peptide partition coefficients spanned more than two orders of magnitude across the library, and RNA enrichment closely tracked peptide enrichment, consistent with co-condensation (Figure 1B). Representative systems illustrate these extremes, with [RRGG]_7_ +RNA forming a clear dense phase and sharply peaked peptide and RNA profiles, whereas [ERGG]_7_ +RNA remained mixed with nearly flat profiles and no distinct dense phase (Figure 1C–F). These simulations indicate that RNA does not act as a generic driver for all repeat peptides; instead, it amplifies intrinsic sequence biases at the X position into pronounced differences in RNA-driven co-phase separation.

**Figure 1.**
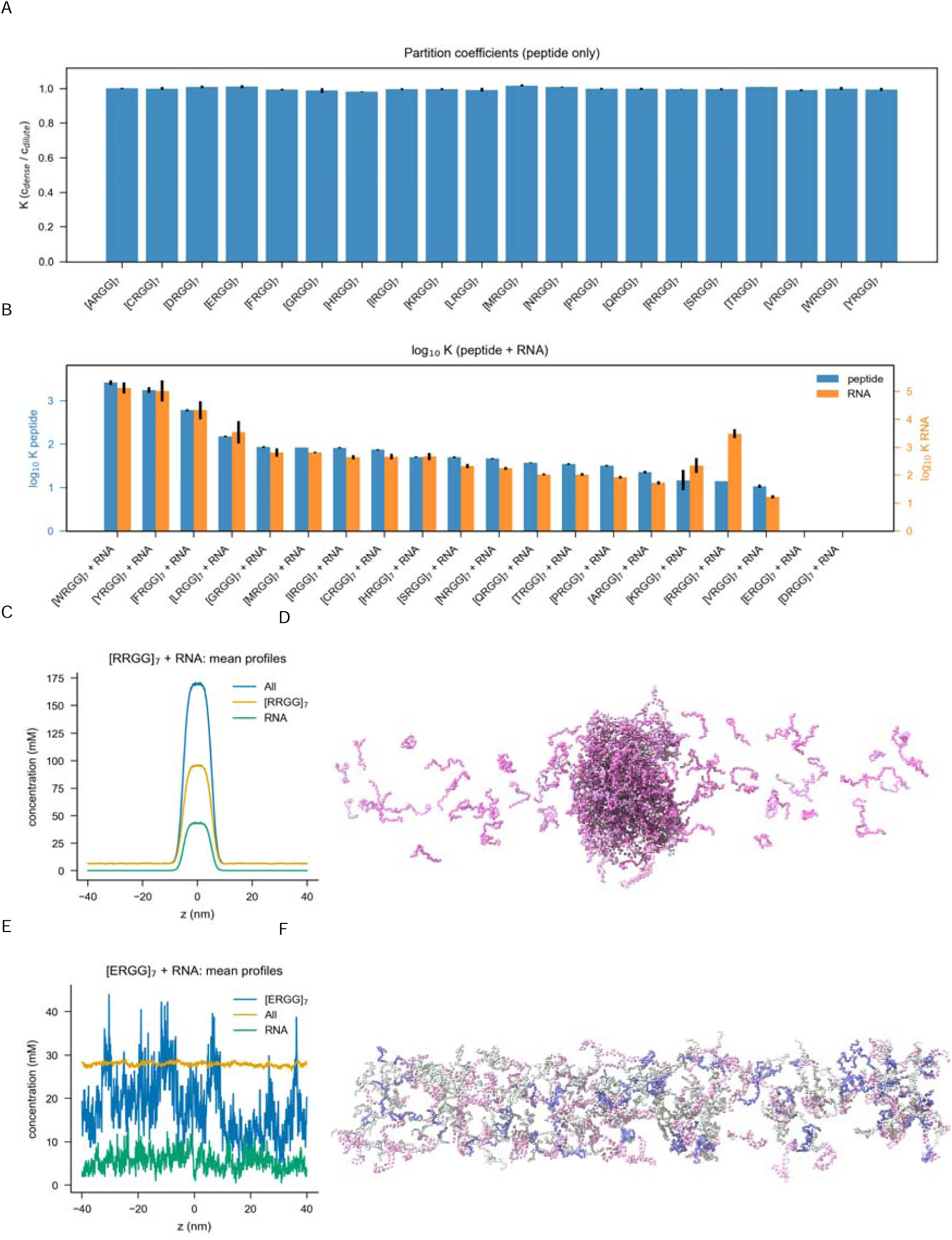
| Coarse-grained simulations reveal sequence-dependent RNA-driven phase separation of [XRGG] peptides. (A) Partition coefficients Kfor peptide-only simulations of the [XRGG]_7_ library. All sequences have K≈1, indicating that [XRGG]_7_ peptides alone do not form a robust condensed phase under these conditions. (B) Partition coefficients for peptide–RNA mixtures, plotted as Blue bars indicate peptide partitioning (left y-axis), orange bars indicate RNA partitioning (right y-axis). Different [XRGG]_7_ sequences now span more than two orders of magnitude in K, revealing strong sequence-dependent co-condensation with RNA. (C) Mean concentration profiles for the [RRGG]_7_ +RNA system. The total (“All”, blue), peptide ([RRGG]_7_, orange) and RNA (green) components all display a pronounced central dense slab and low concentrations in the dilute regions, characteristic of phase separation. (D) Representative snapshot from the [RRGG]_7_ +RNA simulation, showing a compact central condensate surrounded by a dilute gas of peptide and RNA chains. (E) Mean concentration profiles for the [ERGG]_7_ +RNA system. Total, peptide and RNA concentrations remain nearly flat along z, indicating the absence of a distinct dense phase and a mixed, non-separating state. (F) Representative snapshot from the [ERGG]_7_ +RNA simulation, illustrating a laterally extended, low-density distribution of chains without formation of a condensed slab.

Moreover, RG/RGG regions contain variable numbers of RG repeats spaced with different “linker” lengths. To determine the biologically relevant number of RG/RGG repeats in the human IDR dataset (Supplementary file 1), we searched for different numbers of RG/RGG repeats with spacing from zero to five (Supplementary file 4) (41). We found that more than 93% of proteins with RG regions and 85% of proteins with RGG regions contain seven or less repeats (Figure 1A). Therefore, we designed peptides with seven repeats of RGG and all possible preceding residues (termed as a [XRGG]_7_), to study the effect of RG’s preceding residues on phase separation.

RNA can promote phase separation of several RBPs containing RG/RGG regions (50). We therefore monitored condensate formation in a setting with a fixed concentration of [XRGG]_7_ peptides and increasing concentrations of non-specific [UG]_12_ RNA using a turbidity assay that measures the optical density (OD) of a protein solution. [XRGG]_7_ peptides with the following residues in the X position showed increased OD values with increasing concentrations of RNA, indicative of condensate formation: (i) positively charged residues ([RRGG]_7_, [KRGG]_7,_ [HRGG]_7_), (ii) aromatic residue ([YRGG]_7_), (iii) hydrophobic residues ([MRGG]_7_, [LRGG]_7_), (iv) polar residues ([CRGG]_7,_ [NRGG]_7_, [SRGG]_7_), and (v) [GRGG]_7._ In contrast, we observed no or only minor increase in turbidity upon RNA addition for all the other tested peptides (Figure 2B-H).

**Figure 2:**
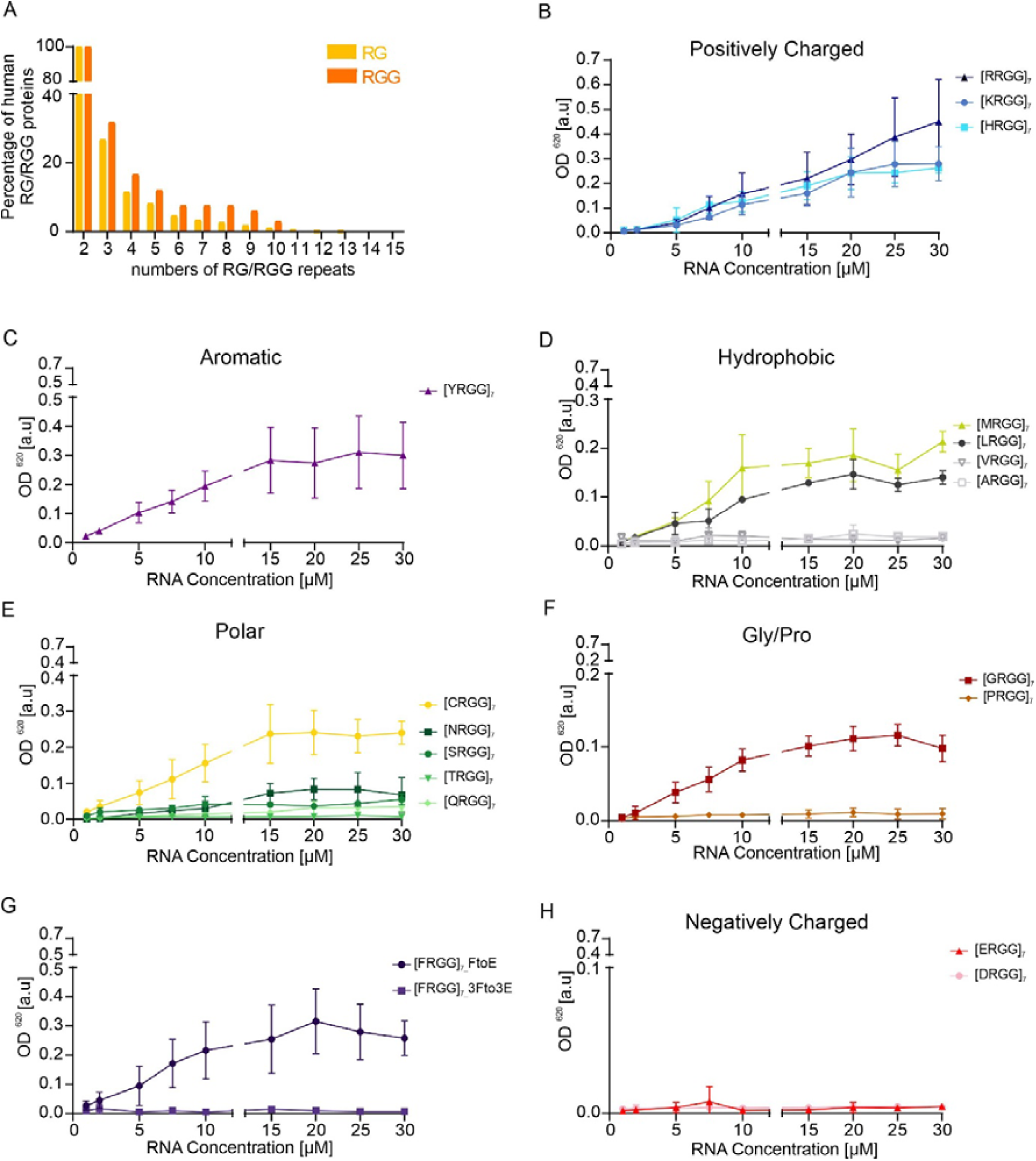
Short RG/RGG repeats are highly abundant in human IDRs and RNA induces condensate formation of [XRGG]_7_ model peptides dependent on the ‘X’ residue. (A) Bioinformatics analysis performed on the abundance of RG or RGG motifs in human IDR dataset, ranging from two to fifteen repeats. The motifs are spaced by zero to five residues (Supplementary file 4). A total number of RG/RGG proteins is calculated from the search for two repeats of RG/RGG with spacing from zero to five. This indicates that 100% of RG/RGG proteins according to our list have at least two repeats of RG/RGG motif. (B-H) Turbidity assays to quantify phase separation of different [XRGG]_7_ model peptides. Experiments were performed with a fixed concentration of peptides (30 µM) and increasing RNA concentration. Values represent means ± SD (n = 3).

To validate our findings, we performed differential interference contrast (DIC) microscopy (Supplementary Figure S1A and B). In the presence of substoichiometric amounts of RNA, the [XRGG]_7_ peptides which showed increased turbidity upon RNA addition immediately formed small condensates that fused and grew in size over time (data not shown) (Supplementary Figure S1A and B). Even though the [SRGG]_7_ peptide did not show any significant increase in turbidity (Figure 2E), we detected formation of small condensates upon RNA addition by DIC (Supplementary Figure S1A). Furthermore, [FRGG]_7_, [WRGG]_7_, and [IRGG]_7_ peptides were prone to form amorphous, irregularly shaped condensates / aggregates both with and without RNA (Supplementary Figure S1A and B). In particular, the addition of RNA to [WRGG]_7_, reduced the formation of such aggregate and shifted morphology toward the formation of smaller droplets (Supplementary Figure S1B). To test if [FRGG]_7_ aggregate formation can be reduced by mutations, we introduced either one negatively charged glutamate in the 4^th^ X position of [FRGG] ([FRGG]_7__FtoE) or three glutamates in the 2^nd^, 4^th^ and 6^th^ X position of [FRGG] ([FRGG]_7__3Fto3E) and tested them in our assays. Interestingly, both modifications lead to solubilization of the aggregates. [FRGG]_7__FtoE phase separated in the presence of RNA whereas [FRGG]_7__3Fto3E did not (Figure 2G).

When comparing the condensate formation properties of the remaining hydrophobic residues in the [XRGG]_7_ background, we found that peptides enriched in hydrophobic residues with long side chains such as [LRGG]_7_ and [MRGG]_7_ can phase separate, whereas peptides with short side chains such as [ARGG]_7_ or [VRGG]_7_ did not form condensates (Figure 2D). Introduction of amino acids strongly affecting the backbone flexibility showed opposite effects, with proline preventing condensate formation and glycine allowing condensate formation (Figure 2F). The peptides enriched in some polar residues, such as [TRGG]_7_ and [QRGG]_7_ did not show RNA-induced condensation by turbidity under these conditions (Figure 2E). These results suggest that positively charged residues and long-chain hydrophobics promote RNA-driven condensation in the [XRGG]_7_ context, while aromatic residues can either support droplet formation (Y) or drive irregular assemblies (F/W/I), depending on sequence context and interaction strength. This data indicates that the sequence context of RG/RGG regions could be a crucial factor in determining phase separation of RG/RGG-containing proteins.

### RG/RGG repeats form compact tracts with short linkers in disordered regions

The [XRGG]_7_ simulations and experiments showed that a single residue immediately preceding RG can tune whether an RG-rich peptide forms RNA-driven condensates. We next asked how RG motifs are arranged in naturally occurring disordered regions of the proteome, and whether there is a characteristic “RG-tract grammar” that could underlie this behaviour.

We first quantified how many RG motifs are present per protein in predicted intrinsically disordered regions (IDRs) and in a curated set of canonical RG/RGG proteins (7,38) (Supplementary File 1). Across all disordered proteins, most sequences contained only a few RG motifs. Proteins from the canonical RG/RGG set were clearly enriched in RGs, but the distribution still peaked below ∼15 motifs per protein (Figure 3A). Thus, even in RG/RGG proteins, RG motifs are not spread uniformly along the chain; instead, they are likely concentrated into local stretches.

To characterise the spacing within these stretches, we next examined the distribution of inter-RG linker lengths (number of residues between consecutive RG motifs). In the disordered proteome, linker lengths spanned a broad range up to ∼25 residues, with substantial weight at intermediate and long linkers (Figure 3B, grey). Canonical RG/RGG proteins, by contrast, were strongly biased towards short linkers: most inter-RG distances were ≤7 residues, with a sharp peak at 0–2 residues and comparatively few long linkers (Figure 3B, blue). These global distributions already suggest that RG/RGG proteins are enriched for compact, tightly spaced RG segments relative to disordered proteins in general.

**Figure 3.**
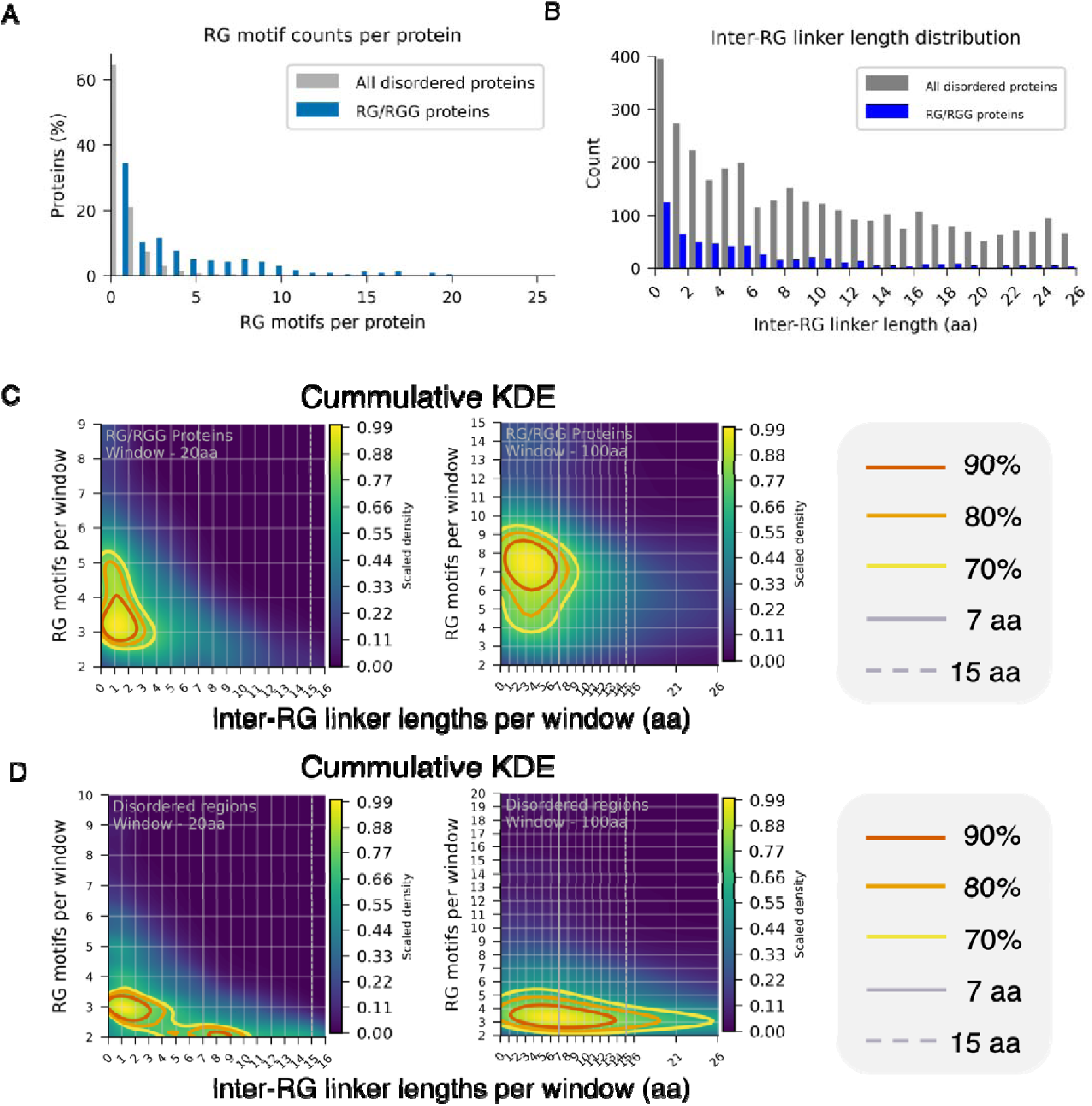
| RG/RGG repeats are enriched in compact tracts with short linkers in disordered regions. (A) Distribution of RG motif counts per protein across all predicted disordered proteins (grey) and the curated RG/RGG protein set (blue). RG/RGG proteins are enriched in RG motifs but still peak below ∼15 motifs per protein, indicating that RG motifs are abundant yet concentrated into local regions rather than uniformly distributed along the sequence. (B) Histogram of inter-RG linker lengths (number of residues between consecutive RG motifs) for all disordered proteins (grey) and RG/RGG proteins (blue). RG/RGG proteins show a strong bias towards short linkers (0–7 residues), whereas disordered proteins in general more frequently harbour longer spacers. (C) Cumulative two-dimensional kernel density estimates (KDEs) of RG organisation in RG/RGG proteins. Each point corresponds to a sequence window containing at least two RG motifs and is described by the number of RG motifs per window (y-axis) and the mean inter-RG linker length (x-axis). (C-left) 20-residue windows; (C-right) 100-residue windows. Contours indicate the 70%, 80% and 90% highest-density regions. Vertical guide lines at 7 residues (solid grey) and 15 residues (dashed grey) mark the compact and permissive linker cutoffs, respectively. Most RG/RGG windows lie to the left of the 7-residue line, consistent with densely packed RG tracts. (D) As in (C), but for all predicted disordered regions in the human proteome. (D-left) 20-residue windows; (D-right) 100-residue windows. Short linkers are still common, but windows with both high RG motif counts and very short mean linkers are comparatively rare, indicating that compact, RG-dense tracts are a distinctive feature of RG/RGG proteins rather than a generic property of disordered regions.

We then asked how this organisation looks in local sequence windows. For each canonical RG/RGG protein, we scanned 20, 40, 60, 80 and 100-residue windows and, for windows containing at least two RG motifs, recorded both the number of RG motifs and the mean inter-RG linker length (contour bounds for all window sizes are provided in Supplementary File 5). Cumulative two-dimensional kernel density estimates (KDEs) revealed that RG motifs in canonical RG/RGG proteins predominantly occupy a “compact” regime (Figure 3C; Supplementary File 5). In short windows (20,40 aa), the 70–90% highest-density contours were centred at 2–4 RG motifs per window and mean linker lengths ≤4 residues (Figure 3C-left). In longer windows (60,80,100 aa), the density shifted towards more motifs per window, but remained largely confined to mean linkers ≤7 residues (Figure 3C-right). Vertical guide lines at 7 and 15 residues highlight that, in RG/RGG proteins, the vast majority of local RG segments fall to the left of the 7-residue cutoff, with only a minority sampling longer spacers.

Applying the same analysis to all predicted IDRs revealed a different pattern. Short disordered windows (20-40 aa) still favoured relatively short linkers, but typically contained fewer RG motifs and showed much less density in the high-motif/high-density regime seen for canonical RG/RGG proteins (Figure 3D-left). In longer windows (60,80,100-residue windows), most disordered regions had mean linkers ≤15 residues, yet only a small subset combined high motif counts with short linkers (Figure 3D-right). In other words, short inter-RG spacers are common across IDRs, but long, densely packed RG tracts are strongly enriched in canonical RG/RGG proteins and comparatively rare elsewhere in the disordered proteome.

Guided by these observations, we defined an operational rule to identify compact RG regions: consecutive RG motifs are assigned to the same tract if separated by ≤7 residues, with a more permissive upper limit of 15 residues used only to bridge otherwise compact segments. Each resulting region was further extended by five residues on both termini to preserve local sequence context. Applying this rule to the whole proteome yielded 1,008 compact RG tracts in 823 proteins (Supplementary File 6), including 380 tracts in 289 annotated RNA-binding proteins. Together with the [XRGG]_7_ data, these analyses support a simple RG-tract grammar: RG motifs are typically arranged in short, tightly spaced clusters with ≤7-residue linkers, and such clusters are particularly enriched in proteins that rely on RG/RGG regions for RNA binding and condensate formation.

### X residues determine TNPO1 binding of XRGG peptides

Some of TNPO1 cargo proteins contain RG/RGG regions (40,41).(25,26,28,31) Specifically, a proteomics screen identified 72 out of 643 TNPO1 cargoes harboring RG/RGG regions (Figure 4A and Supplementary files 4-6), suggesting that their interaction through RG/RGG regions is important for their nuclear import. Although RG/RGG regions were identified several years ago to be involved in TNPO1-mediated nuclear import (25,26,28,31), the primary sequence determinants regulating the interaction between RG/RGG regions and TNPO1 are yet not well understood. Here, we hypothesized that the residues surrounding the RG/RGG regions are key in TNPO1 recognition. To further investigate this, we determined the TNPO1 binding affinities of [XRGG]_7_ peptides using isothermal titration calorimetry (ITC) (Table 1; Figure 4B; and Supplementary Figure S2). [XRGG]_7_ peptides with positively charged residues at the [X] position exhibited the highest TNPO1 binding affinities in the nanomolar (nM) K_D_ range. Specifically, [KRGG]_7_ and [HRGG]_7_ showed affinities of 150 ± 30 nM and 770 ± 70 nM, respectively (Table 1; Figure 4B; and Supplementary Figure S2). On the other hand, [RRGG]_7_ exhibited a different binding mode, with two binding sites on TNPO1, with affinities of 207 ± 72 pM and 135 ± 10 nM for the first and second site, respectively (Supplementary Figure S2, and Table 1). Among the peptides with aromatic X residues, and due to the aggregate formation observed for [FRGG]_7_ and [WRGG]_7_, we could determine binding affinities only for [YRGG]_7_. Interestingly, ITC revealed that TNPO1 can bind two molecules of [YRGG]_7_ (Supplementary Figure S2), with an affinity in the high nanomolar range (700 ± 90 nM) (Table 1, and Figure 4B). The [FRGG]_7_ peptides harbouring either one glutamate substitution ([FRGG]_7__FtoE) or three glutamate substitutions ([FRGG]_7__3Fto3E) displayed TNPO1 binding affinities of 430 ± 60 nM and 1.5 ± 0.5 µM, respectively (Supplementary Figure S2; Table 1). [XRGG]_7_ peptides with hydrophobic amino acids with long side chains (Met or Leu) in the X position displayed high nanomolar binding affinities of 877 ± 53 nM and 790 ± 70 nM, respectively, for TNPO1 (Supplementary Figure S2; Figure 4B, and Table 1). [XRGG]_7_ peptides harboring either hydrophobic amino acids (Val and Ala), polar amino acids (Cys, Asn, Ser, Thr and Gln), Gly, or Pro in the X position, showed weaker binding to TNPO1 with K_D_s ranging from 1.2 µM to 2.3 µM (Supplementary Figure S2; table 1, and Figure 4B). [ERGG]_7_ and [DRGG]_7_ containing negatively charged residues at the X position did not show detectable binding to TNPO1 (Supplementary Figure S2; table 1, and Figure 4B). In summary, we found that the X residue in an [XRGG]_7_ sequence can alter TNPO1 binding and that [XRGG]_7_ peptides with a high propensity for condensate formation displayed the highest binding affinities for TNPO1, whereas [XRGG]_7_ peptides not undergoing phase separation showed weaker or no binding to TNPO1 (Figure 4C). This correlation guided us to investigate the chaperoning function of TNPO1 on peptides that undergo phase separation.

**Figure 4:**
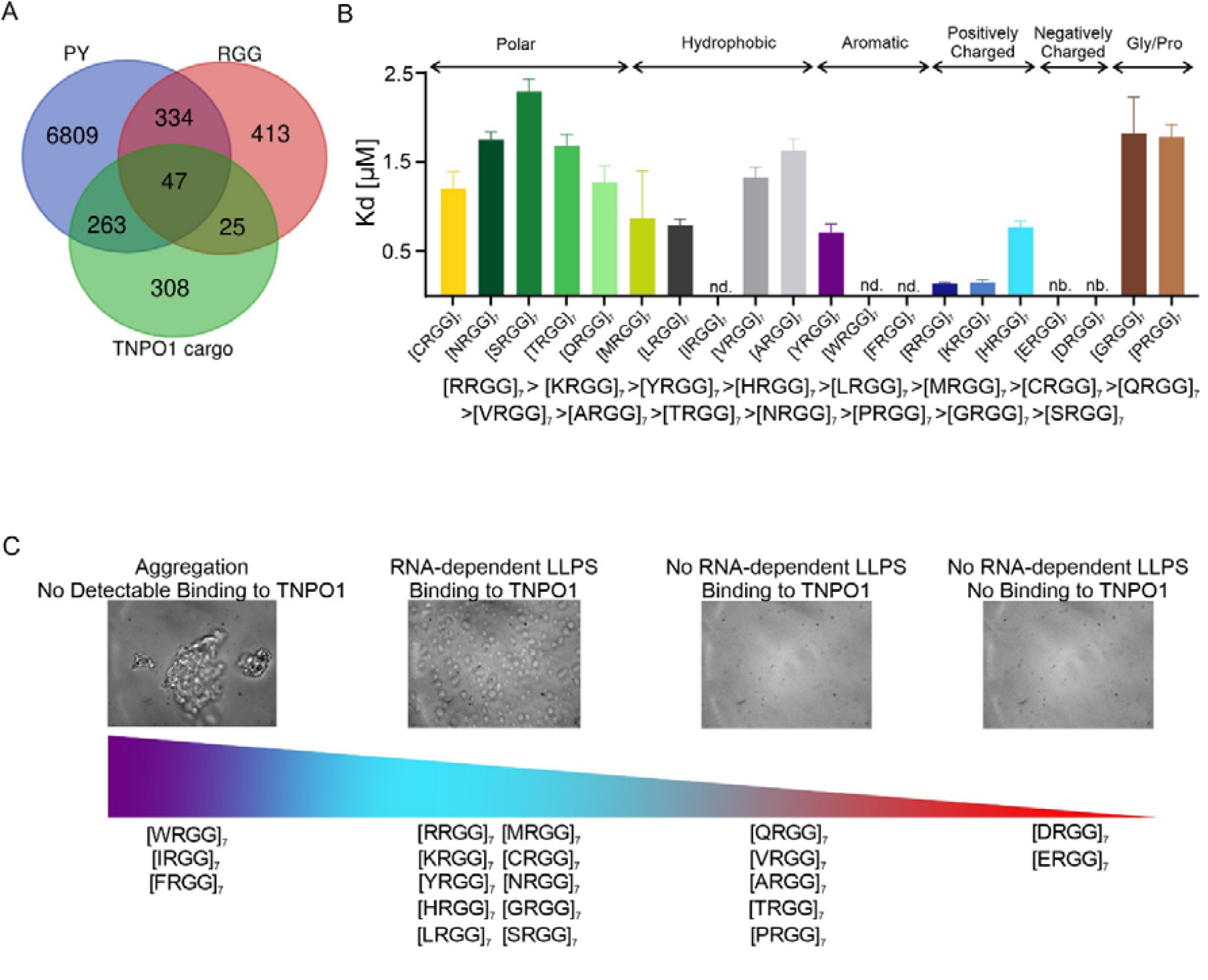
RG/RGG proteins are TNPO1 cargoes and [XRGG]_7_ model peptides have affinities (K_D_) within nanomolar to low micromolar range for TNPO1. (A) Venn diagram corresponding to the PROSITE analysis (https://prosite.expasy.org/scanprosite/) of the PY motif (blue) preceded by a basic amino acid spaced by zero to nine residues [[RKH]-x(0–9)-P-Y] within human proteins (taxid:9606) (Supplementary file 7) and the selected RG-rich regions (Supplementary file 6) within human IDRs dataset (Supplementary file 1). These two motifs were compared with TNPO1 cargoes (green) which were identified and published by Kimura *et al.* (40) and Mackmull *et al.* (41) (Supplementary file 7). (B) K_D_ values of [XRGG]_7_ peptides for TNPO1 measured by titration of 100 µM peptides into 10 µM of TNPO1. 200 µM [YRGG]_7_ was titrated into 10 µM TNPO1 because two molecules of [YRGG]_7_ bind to one molecule of TNPO1. 50 µM [FRGG]_7_ was titrated into 5 µM TNPO1 because [FRGG]_7_ was not soluble at higher concentrations. (nd: not detectable, nb: no binding). No binding was detectable for [DRGG]_7_ and [ERGG]_7,_ and affinities of [WRGG]_7_, [IRGG]_7_ and [FRGG]_7_ to TNPO1 are not detectable. (C) Schematic summary of peptides categorized according to their ability to phase separate and to bind to TNPO1.

**Table 1:** Thermodynamic parameters of ITC titrations. The reported errors correspond to the SD of the fit. (n.b: no detectable binding, n.d. not detectable)

| Cell | Syringe | K<sub>d</sub> (μM) | ΔH (kcal·mol<sup>-1</sup>) | ΔS (cal/mol·K) |
| --- | --- | --- | --- | --- |
| **TNPO1** | [GRGG]<sub>7</sub> | 1.82 ± 0.41 | -4.00 ± 0.24 | 12.8 |
| **TNPO1** | [ERGG]<sub>7</sub> | n.b. | - | - |
| **TNPO1** | [YRGG]<sub>7</sub> | 0.70 ± 0.09 | -8.54 ± 0.16 | -0.47 |
| **TNPO1** | [FRGG]<sub>7</sub> | n.d. | - | - |
| **TNPO1** | [ARGG]<sub>7</sub> | 1.63 ± 0.13 | -7.39 ± 0.18 | 1.69 |
| **TNPO1** | [VRGG]<sub>7</sub> | 1.33 ± 0.11 | -6.70 ± 0.17 | 4.43 |
| **TNPO1** | [CRGG]<sub>7</sub> | 1.20 ± 0.19 | -8.34 ± 0.27 | -0.88 |
| **TNPO1** | [PRGG]<sub>7</sub> | 1.78 ± 0.14 | -6.48 ± 0.16 | 4.58 |
| **TNPO1** | [LRGG]<sub>7</sub> (240s spacing) | 0.79 ± 0.07 | -5.98 ± 0.11 | 7.87 |
| **TNPO1** | [IRGG]<sub>7</sub> | n.d. | - | - |
| **TNPO1** | [MRGG]<sub>7</sub> | 0.87 ± 0.053 | -9.57 ± 0.12 | -4.38 |
| **TNPO1** | [WRGG]<sub>7</sub> | n.d. | - | - |
| **TNPO1** | [KRGG]<sub>7</sub> | 0.15 ± 0.03 | -7.88 ± 0.18 | 4.79 |
| **TNPO1** | [HRGG]<sub>7</sub> | 0.77 ± 0.07 | -6.36 ± 0.11 | 6.64 |
| **TNPO1** | [SRGG]<sub>7</sub> | 2.29 ± 0.14 | -12.70 ± 0.34 | -16.8 |
| **TNPO1** | [TRGG]<sub>7</sub> | 1.68 ± 0.13 | -7.68 ± 0.21 | 0.68 |
| **TNPO1** | [NRGG]<sub>7</sub> | 1.75 ± 0.09 | -10.30 ± 0.17 | -8.22 |
| **TNPO1** | [QRGG]<sub>7</sub> | 1.27 ± 0.19 | -7.62 ± 0.30 | 1.43 |
| **TNPO1** | [DRGG]<sub>7</sub> | n.b. | - | - |
| **TNPO1** | [FRGG]<sub>7</sub>FtoE | 0.43 ± 0.06 | -4.45 ± 0.07 | 14.2 |
| **TNPO1** | [FRGG]<sub>7</sub>3Fto3E | 1.48 ± 0.46 | -3.42 ± 0.32 | 15.2 |
| **TNPO1** | [ERGG]<sub>7</sub>EtoF | n.b. | - | - |
| **TNPO1** | [ERGG]<sub>7</sub>3Eto3F | 1.04 ± 0.40 | -1.90 ± 0.15 | 21.0 |
|  |  | **K<sub>d</sub> (pM – nM)** |  |  |
| **TNPO1** | [RRGG]<sub>7</sub> | K<sub>d1</sub>: 207 ± 72 pM<br>K<sub>d2</sub>: 135 ± 10 nM | -14.05 ± 0.06<br>-10.49 ± 0.09 | -2.82<br>-3.75 |
|  |  | **K<sub>d</sub> (μM)** |  |  |
| **TNPO1** | HNRPU<sup>700-730</sup> | 1.55 ± 0.23 | -4.50 ± 0.26 | 11.5 |
| **TNPO1** | HNRL1<sup>637-659</sup> | n.b. | - | - |
| **TNPO1** | NUCL<sup>653-696</sup> | 0.30 ± 0.05 | -9.89 ± 0.20 | -3.32 |
| **TNPO1** | FBRL<sup>8-47</sup> | 0.40 ± 0.04 | -9.13 ± 0.13 | -1.35 |
| **TNPO1** | LS14A<sup>401-431</sup> | 0.67 ± 0.06 | -13.30 ± 0.24 | -16.3 |
| **TNPO1** | PAIRB<sup>363-387</sup> | 0.40 ± 0.04 | -11.16 ± 0.18 | -8.16 |
|  |  | **Kd (nM)** |  |  |
| **TNPO1** | FUS<sup>RGG3-PY</sup> (240s spacing) | 36 ± 9 | -33.6 ± 0.51 | -78.5 |
| **TNPO1** | X6C_ FUS<sup>RGG3-PY</sup> | 41 ± 5 | -9.95 ± 0.09 | 0.4 |
| **TNPO1** | X6E_ FUS<sup>RGG3-PY</sup> | 32 ± 5 | -31.3 ± 0.34 | -70.6 |
| **TNPO1** | X6T_ FUS<sup>RGG3-PY</sup> | 28 ± 5 | -29.5 ± 0.36 | -64.5 |
| **TNPO1** | X6R_ FUS<sup>RGG3-PY</sup> (240s spacing) | 164 ± 27 | -24.9 ± 0.60 | -52.5 |
| **TNPO1** | X6Y_ FUS<sup>RGG3-PY</sup> | 44 ± 11 | -17.7 ± 0.32 | -25.6 |
| **TNPO1** | X6F_ FUS<sup>RGG3-PY</sup> | 36 ± 12 | -19.8 ± 0.47 | -32.4 |

Previous studies showed that TNPO1 acts as a molecular chaperone for RBPs with low complexity sequences, including CIRBP and FUS (28,30–32). Therefore, we tested how TNPO1 affects RNA-mediated [XRGG]_7_ peptide condensate formation observed here using turbidity assays and DIC microscopy. The addition of increasing amounts of TNPO1 resulted in a concentration-dependent decrease in turbidity of the tested [XRGG]_7_ peptides, indicating that TNPO1 can completely inhibit phase separation of [CRGG]_7_, [GRGG]_7_, [MRGG]_7_, [LRGG]_7_, and [NRGG]_7_ (Figure 5A and E). However,

**Figure 5:**
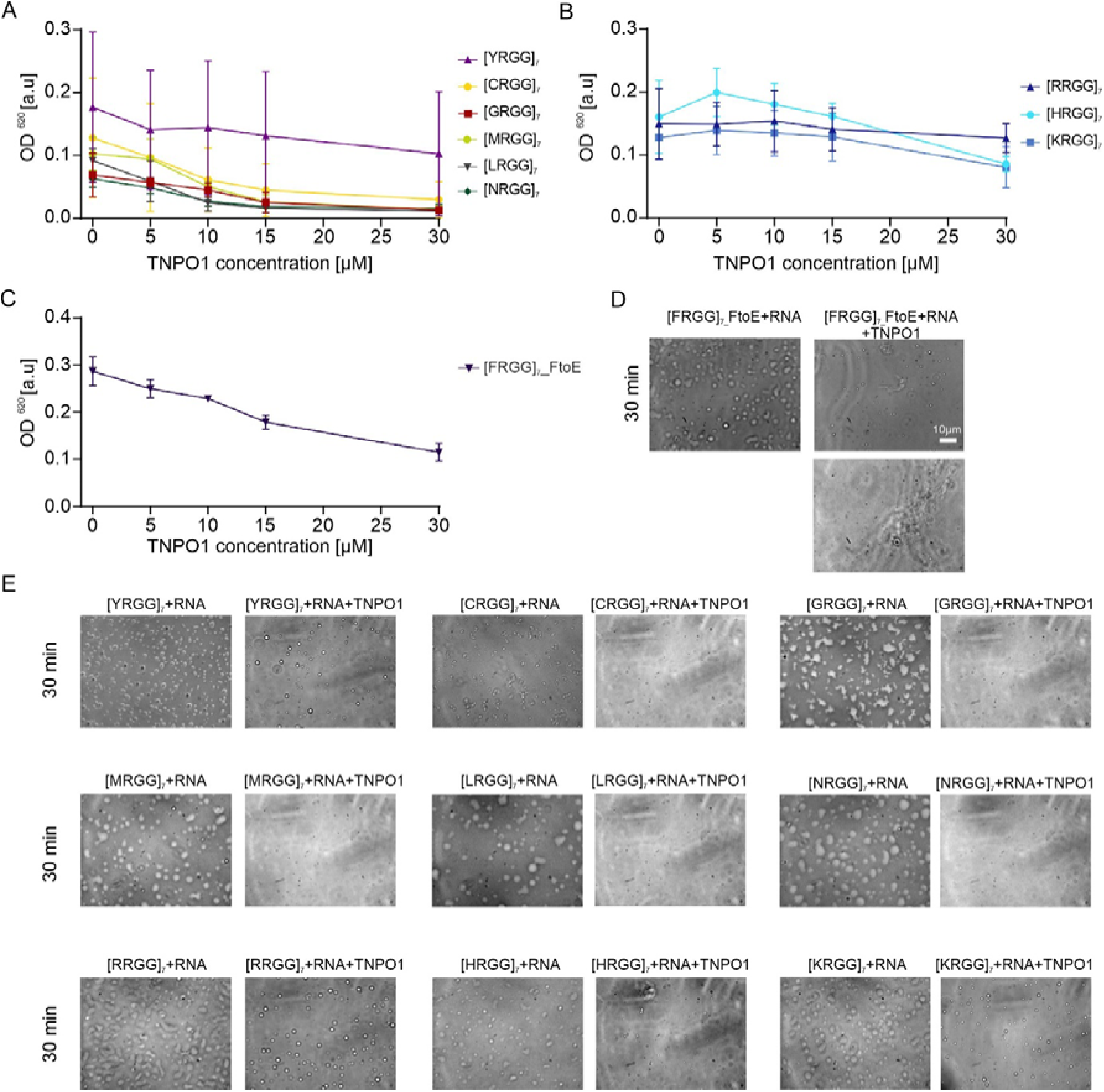
Chaperoning of [XRGG]_7_ peptides by TNPO1. (A-C) Turbidity assays to quantify phase separation of peptides with a fixed concentration of peptides (30 µM) and RNA (10 µM) and increasing concentration of TNPO1 measured at OD 620 nm immediately after sample preparation. Values represent means ± SD (n = 3). (D-E) Differential interference contrast microscope images of 30 µM [XRGG]_7_ peptides in the presence of 15 µM RNA or 10 µM RNA with 30 µM TNPO1 after 30 min. (Scale bar: 10µm)

TNPO1 only partially inhibits the phase separation of [YRGG]_7_, [HRGG]_7_, [RRGG]_7_, [KRGG]_7_ and [FRGG]_7__FtoE (Figure 5A-E). In agreement, a strong impairment of condensate formation of the [XRGG]_7_ peptides was observed in the presence of TNPO1 by DIC microscopy (Figure 5D and E). As a summary, we demonstrated that TNPO1 can inhibit phase separation of [XRGG]_7_ peptides.

### Sequence context determines condensate formation and TNPO1-binding

To further investigate the impact of sequence context in human RG/RGG regions beyond our simple model peptides, we used the PROSITE database (https://prosite.expasy.org/scanprosite), which enables pattern-based searches of protein sequences. We identified RG/RGG-containing proteins with putatively high propensity to phase separate and to interact with TNPO1, based on the presence of Met, Tyr, Lys, His, Arg, Leu, Gly, or Phe at the [X] position. The selected RBPs are shown in Figure 4A, and include heterogeneous nuclear ribonucleoprotein U (HNRPU^700-730^), heterogeneous nuclear ribonucleoprotein U-like protein 1 (HNRL1^637-659^), nucleolin (NUCL^653-695^), rRNA 2’-O-methyltransferase fibrillarin (FBRL^8-47^), protein LSM14 homolog A (LS14A^401-431^) and plasminogen activator inhibitor 1 RNA-binding protein (PAIRB^363-387^) (Figure 6A). We therefore examined whether these natural RG/RGG regions indeed have a high propensity to phase separate, to bind to TNPO1 and to be chaperoned by TNPO1, which is what we would predict based on our experiments carried out with the [XRGG]_7_ model peptides.

**Figure 6:**
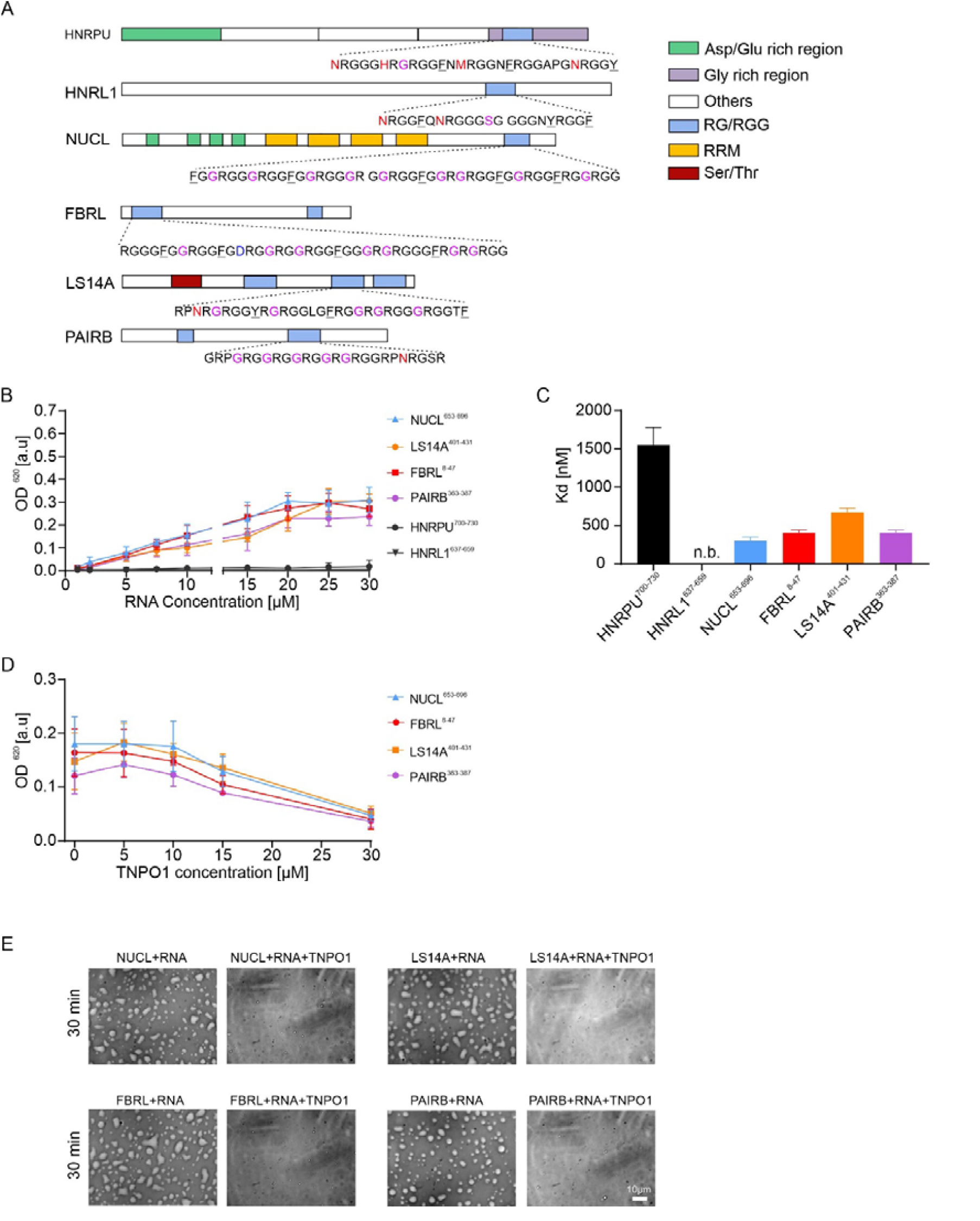
Natural human RG/RGG-containing peptides with high net charge and enriched by aromatic residues undergo phase separation, bind TNPO1, and can be chaperoned by TNPO1. (A) Schematic diagrams of chosen candidate proteins and their RG/RGG sequences. (B) Quantification for phase separation of natural human RG/RGG-containing sequences at a fixed concentration of peptide (30 µM) and increasing concentration of RNA. Values represent means ± SD (n = 3). (C) Graph shows K_D_ values of RG/RGG containing natural human peptides for TNPO1, determined by titration of 100 µM peptides into 10 µM of TNPO1 with ITC. (D) Quantification for phase separation of RG/RGG-containing natural human proteins at a fixed concentration of peptide (30 µM) and RNA (10 µM) and increasing concentration of TNPO1. Values represent means ± SD (n = 3). (E) DIC microscope images of 30 µM RG/RGG-containing human peptides in the presence of 15 µM RNA or 10 µM RNA with 30 µM TNPO1 after 30 min. (Scale bar: 10 µm)

Indeed, natural RG/RGG regions with a high net charge (around 9 to10), such as NUCL^653-695^, FBRL^8-47^, LS14A^401-431^, and PAIRB^363-387^ (Figure 6A and Table 2) underwent phase separation, as observed by increased turbidity and formation of condensates (Figure 6B and 6E). However, RG/RGG regions with a low net charge (less than 7), such as HNRPU^700-730^ and HNRL1^637-659^, did not undergo phase separation (Figure 6A-B and Table 2), despite containing His, Met, and Tyr residues that allow for phase separation in the [XRGG]_7_ context (Figure 2B,D and E and Figure 6A-B). Thus, we conclude that in addition to the X residue, high net charge is a key determinant of phase separation of natural RG/RGG peptides.

**Table 2:**
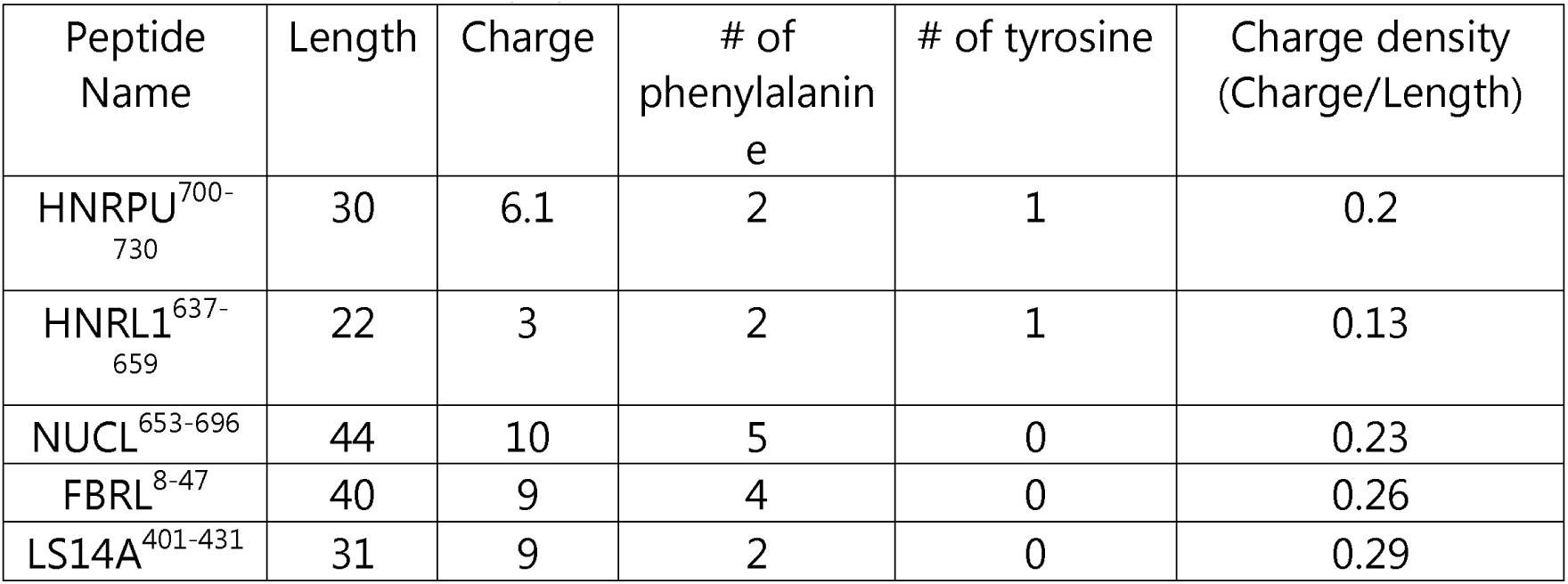
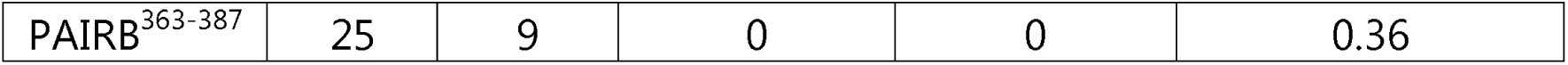
Information about peptide features.

We then assessed whether a correlation exists between the propensity of RG/RGG regions to phase separate and their ability to interact with TNPO1, as observed for the [XRGG]_7_ peptides. In ITC measurements we observed that RG/RGG regions with high net charge, such as NUCL^653-695^, FBRL^8-47^, LS14A^401-431^ and PAIRB^363-387^ interacted with TNPO1 with an associated K_D_ in the nanomolar range, of 300 ± 50 nM, 400 ± 40 nM, 670 ± 60 nM, and 400 ± 40 nM, respectively (Figure 6C; Supplementary Figure S3C-F; and Table 1). The affinity of HNRPU^700-730^ for TNPO1 was lower (1.55 ± 0.23 µM) and HNRL1^637-659^, which also contains an RG/RGG region with a low net charge, did not bind to TNPO1 at all (Figure 6C; Supplementary Figure S3C-F; and Table 1). This data suggests that the charge density (charge/length) of the protein sequence maybe an important determinant for TNPO1 binding (Table 2).

Finally, we tested the “chaperoning” function of TNPO1 with respect to condensate formation of RG/RGG-containing proteins. In line with our expectations, addition of increasing amounts of TNPO1 resulted in a concentration-dependent decrease in turbidity of the tested RG/RGG regions, indicating that TNPO1 inhibits their phase separation (Figure 6D). Further confirming these results, impairment of condensate formation was observed in the presence of TNPO1 by DIC microscopy (Figure 6E).

Taken together, using RG/RGG sequences from natural proteins, we show that sequence context modulates RNA-driven condensation and TNPO1 binding *in vitro*, with net charge and charge density contributing strongly under our assay conditions. However, SG recruitment is a complex, multi-component cellular phenomenon and is therefore not expected to directly mirror these *in vitro* readouts; Fig. 8 provides an orthogonal cellular context, and discrepancies likely reflect additional determinants of SG partitioning.

### Mutational studies using FUS reveal neutral and dominant effects

To validate our findings obtained for natural RG/RGG regions in the context of an entire protein, we chose the representative RG/RGG protein FUS as a model system. FUS is a nucleic acid binding protein that is predominantly localized in the nucleus (51–54), and has roles in various nuclear processes like transcription, splicing and microRNA processing (55,56). FUS harbors a N-terminal low-complexity domain and three RG/RGG regions which are important for phase separation (25,57). Mutations in FUS result in the impairment of nuclear localization and the formation of irreversible aggregates in the cytoplasm, which are a hallmark of neurodegenerative disease (31,45,47). Interestingly, a few mutations within the RGG region localize in “[X]” positions, such as D480N in kidney carcinoma tissue (58) and S513P, which was found in 40 familial ALS cases and has also been associated with spinal progressive muscular atrophy (59,60).

Nuclear import of FUS is mediated by TNPO1, which recognizes the PY-NLS located in FUS’ far C-terminal region. Previously, we and others have shown that the FUS^RGG3^ region contributes to TNPO1 recognition, and that this is modulated by mutations and arginine methylation (31,45,46). In this study, we therefore first focused on the FUS^RGG3-PY^ region to better understand the role of surrounding residues in phase separation and TNPO1-mediated chaperoning. To this end, we generated FUS^RGG3-PY^ protein variants by substituting residues at positions 472, 480, 486, 490, 494, and 502 either with Arg, Tyr, Phe, Cys, Glu or Thr, respectively (Figure 7A). Based on our data obtained for the RG/RGG model peptides, we expected that introducing Arg, Tyr, Phe, and Cys would increase phase separation, while introducing Glu and Thr would decrease it (Figure 2B-H). In line with this hypothesis, introduction of aromatic residues (Tyr and Phe) into the FUS^RGG3-PY^ region increased phase separation in presence of RNA, while the introduction of positively charged Arg mutations did not affect it (Figure 7B). Therefore, aromatic residues seem to have a stronger impact on phase separation of FUS^RGG3-PY^ than the positively charged residue Arg. Conversely, introduction of Cys and Thr did not significantly affect phase separation. Negatively charged (Glu) mutations abolished condensate formation as expected (Figure 7B). Glu mutations in FUS^RGG3-PY^ changed the net charge of the protein from 7.2 to 3.2 (Table 3), which might interfere with RNA interaction and abolish RNA-dependent phase separation. In summary, introducing aromatic residues (Tyr and Phe) and negatively charged Glu mutations in a natural sequence context strongly tuned the RNA-dependent phase separation of FUS^RGG3-PY^ (dominant), while introducing Arg, Cys and Thr mutations had only weak or no effects (neutral).

**Figure 7:**
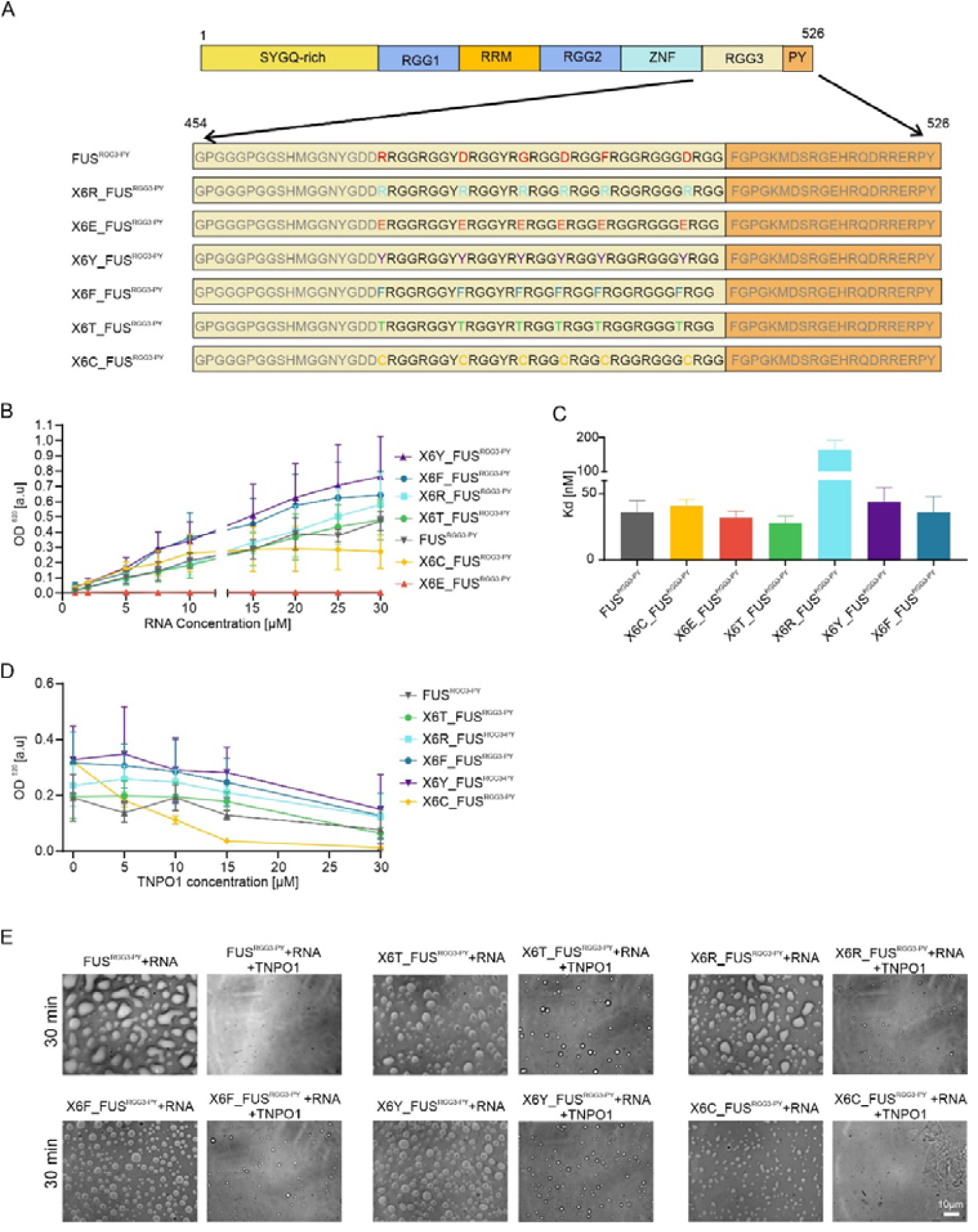
X residues preceding the arginines in the FUS^RGG3-PY^ region regulate phase separation and chaperoning by TNPO1. (A) Schematic diagram of the FUS protein and mutations which were introduced in the FUS^RGG3-PY^region. (B) Quantification of phase separation for WT and the indicated FUS^RGG3-PY^ mutant at a fixed protein concentration (30 µM) and increasing concentration of RNA measured by turbidity at OD 620 nm immediately after sample preparation. Values represent means ± SD (n = 3). (C) Graph shows K_D_ values of WT and mutated FUS^RGG3-PY^ for TNPO1 which were found by titration of 50 µM proteins into 5 µM of TNPO1 with ITC. (D) Quantification for phase separation of WT and mutated FUS at a fixed concentration of peptide (30 µM) and RNA (10 µM) and increasing concentration of TNPO1. Values represent means ± SD (n = 3). (E) DIC microscope images of 30 µM WT and mutated FUS in the presence of 15 µM RNA or 10 µM RNA with 30 µM TNPO1 after 30 min. (Scale bar: 10 µm).

**Table 3:** The net charge of FUS^RGG3-PY^ mutants.

| Protein Name | Charge |
| --- | --- |
| FUS<sup>RGG3-PY</sup> | 7.2 |
| X6Y_ FUS<sup>RGG3-PY</sup> | 9.2 |
| X6F_ FUS<sup>RGG3-PY</sup> | 9.2 |
| X6R_ FUS<sup>RGG3-PY</sup> | 15.2 |
| X6T_ FUS<sup>RGG3-PY</sup> | 9.2 |
| X6C_ FUS<sup>RGG3-PY</sup> | 8.8 |
| X6E_ FUS<sup>RGG3-PY</sup> | 3.2 |

We then investigated whether mutations in X residues of the FUS^RGG3-PY^ region affect TNPO1-binding using ITC (Supplementary Figure S4A-G and Figure 7C). Mutated FUS constructs did not show a significant effect on the binding of TNPO1, except for Arg introductions (X6R), which decreased TNPO1 binding affinity by approximately 3-fold, indicating interference with the “classical” PY-NLS binding to TNPO1 (Supplementary Figure S4C and Figure 7C). To further evaluate this hypothesis, we conducted a competition assay using fluorescence polarization and observed that [RRGG]_7_ interfered with the binding of PY-NLS to TNPO1 (Supplementary Figure S5). We used a higher concentration of reducing reagent compared to previous assays and other mutated version to prevent the formation of disulfide bonds between cysteines in X6C_FUS^RGG3PY^ and found that two molecules of X6C_FUS^RGG3PY^ can bind to TNPO1 (Supplementary Figure S4F). To investigate how the sequence context affects the chaperoning of FUS phase separation by TNPO1, we measured turbidity of the different FUS^RGG3-PY^ mutants under conditions of RNA-mediated phase separation upon addition of TNPO1. We found that TNPO1 partially inhibited phase separation of all FUS^RGG3-PY^ mutants with the exception of X6C_FUS^RGG3-PY^, whose phase separation was impaired completely by addition of TNPO1 (Figure 7D and E).

Our findings highlight the significant impact of the residues surrounding the arginines in RG/RGG residues in FUS^RGG3-PY^ on phase separation. Specifically, we found that aromatic residues and the net charge of the protein are key determinants for the formation of FUS^RGG3-PY^ condensates, while the introduction of negatively charged residues, such as Glu, can disrupt phase separation. We show that the highly positively charged RGG3 region of FUS can interfere with the binding of TNPO1 to PYNLS. Moreover, we observed that the increased phase separation propensity of FUS mutants slightly decreased the efficiency of TNPO1-chaperoning.

### The sequence context of the RG/RGG region modulates SG localization

To validate our *in vitro* findings in cells, we performed a SG recruitment assay in semi-permeabilized HeLa cells (61) using FITC-labeled RG/RGG regions of human proteins (HNRPU^700-730^, HNRL1^637-659^, NUCL^653-695^, FBRL^8-47^, LS14A^401-431^ and PAIRB^363-387^) and Atto-labeled short FUS^RGG3-PY^ constructs (FUS^471-526^, X6E FUS^471-526^ and X6R FUS^471-526^). Stress granules were induced prior to semi-permeabilization and peptides were then introduced to test recruitment into these preformed SGs. We used semi-permeabilized cells to introduce defined, fluorescent peptides directly, rather than relying on transfection/expression. Briefly, SG formation was induced by MG132 treatment for 3 hours; cells were then permeabilized by addition of digitonin, and nuclear pores were blocked by WGA to prevent active nuclear import of peptides. Then, FITC and Atto-labelled peptides were added to the cells and their localization to SGs was monitored. Human CIRBP^89-125^ was used as a positive control ^24^. SGs were visualized by immunostaining for GTPase-activating protein-binding protein (G3BP1).

To relate the intrinsic in vitro condensation and TNPO1-binding propensities to a cellular context, we assessed partitioning into stress granules as an orthogonal readout, noting that SG recruitment is governed by additional determinants beyond the minimal *in vitro* assays. We found that RG/RGG-containing natural peptides were not or less efficiently (NUCL) recruited into preformed SGs compared to our positive control CIRBP and instead mainly diffused through the membrane to the nucleus (Figure 8A-B). Moreover, as expected, we observed that additional arginines (X6R) in the FUS^RGG3-PY^ region significantly increased SG recruitment, whereas introduction of additional glutamines (X6E) decreased SG recruitment of the FUS^RGG3-PY^ protein (Figure 8C-D). In summary, our data suggest that the net charge modulates recruitment into SGs, but the stronger SG colocalization of FUS-RGG-PY despite its lower net charge indicates that additional sequence, such as the PY-NLS/TNPO1-interacting module and other multivalent contacts, also contribute to SG partitioning *in vivo*.

**Figure 8:**
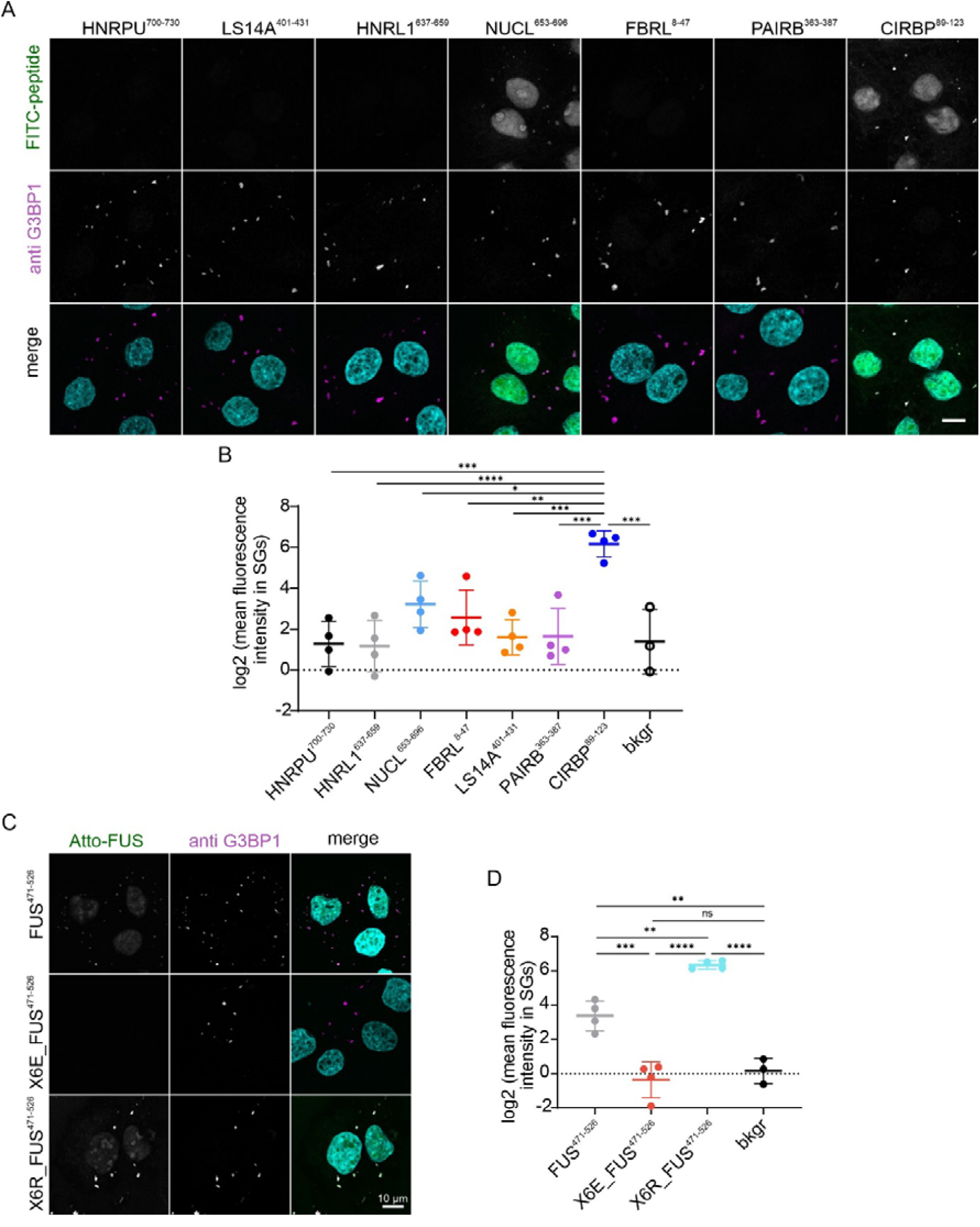
RG/RGG sequences with high net charge are recruited into stress granules. SG recruitment assay using FITC labelled NUCL^653-695^, FBRL^8-47^, LS14A^401-431^, PAIRB^363-387^, HNRPU^700-730^, HNRL1^637-659^ and CIRBP^89-125^ (A, B) or Atto labelled FUS^471-526^, X6E_FUS^471-526^ and X6R_ FUS^471-526^ (C, D) in semi-permeabilized HeLa cells. SGs were visualized by anti-G3BP1 immunostaining, and the recruitment of peptides into SGs was monitored directly by FITC fluorescence using confocal microscopy. Nuclei were counterstained with DAPI. Scale bar, 10 µm. Quantification of fluorescence intensity is shown as the mean ± SD (4 independent replicates) in comparison to a background control (bkgr), where no peptide was added. Statistical significance was calculated by one-way ANOVA with Tukey’s multiple comparison tests (*P<0.05, **P<0.005, ***P<0.0005, ****P<0.0001, ns: not significant).

### Residues surrounding the RG/RGG region modulate nuclear import of FUS

While RG/RGG regions can modulate the nuclear import of RBPs, it remains unclear how these regions mediate the interaction between cargoes and the nuclear import receptor TNPO1. To investigate the role of residues surrounding arginines in the RG/RGG regions and to validate our *in vitro* TNPO1 binding data in intact cells, we included the full-length FUS in our studies. We introduced mutations (Arg, Thr, Phe or Glu) in preceding X residues within all three FUS RGG domains and used our established hormone-inducible import assay to monitor the effect of these mutations on the nuclear import of FUS (62,63). In brief, FUS is fused to two hormone binding domains of the glucocorticoid receptor, which keep the fusion protein in the cytosol after translation, and to two GFP moieties for visualization (GCR_2_-GFP_2_-FUS). Upon addition of the steroid hormone dexamethasone the fusion reporter is released and its nuclear import can be monitored over time by live cell imaging. HeLa cells were transfected with GCR_2_-GFP_2_-FUS or the respective XRGG mutants and their nuclear import rates upon dexamethasone addition were followed over time (Figure 9A-B). Interestingly, we observed that FUS reporters carrying Thr, Arg, Phe or Glu substitutions in the X residue showed a significant decrease compared to the nuclear import of wildtype FUS, despite an intact PY-NLS. Interestingly, Arg and Thr mutants still display some nuclear import when compared to the control, whereas Phe and Glu mutations very strongly inhibited nuclear import. This is in partial agreement with the *in vitro* TNPO1 binding data obtained for (mutant) FUS^RGG3-PY^ peptides (Figure 7C) and suggests that other mechanisms, besides PY-NLS binding, contribute to nuclear import of FUS, such as binding of TNPO1 to the FUS RG/RGG regions and recruitment of FUS to membrane-less organelles (31). Given that a plethora of other RBPs share a similar architecture, such as a PY-NLS in combination with RG/RGG regions, we propose that RG/RGG regions could be crucial in determining nuclear import and intracellular localization of such RG/RGG-containing RBPs.

**Figure 9:**
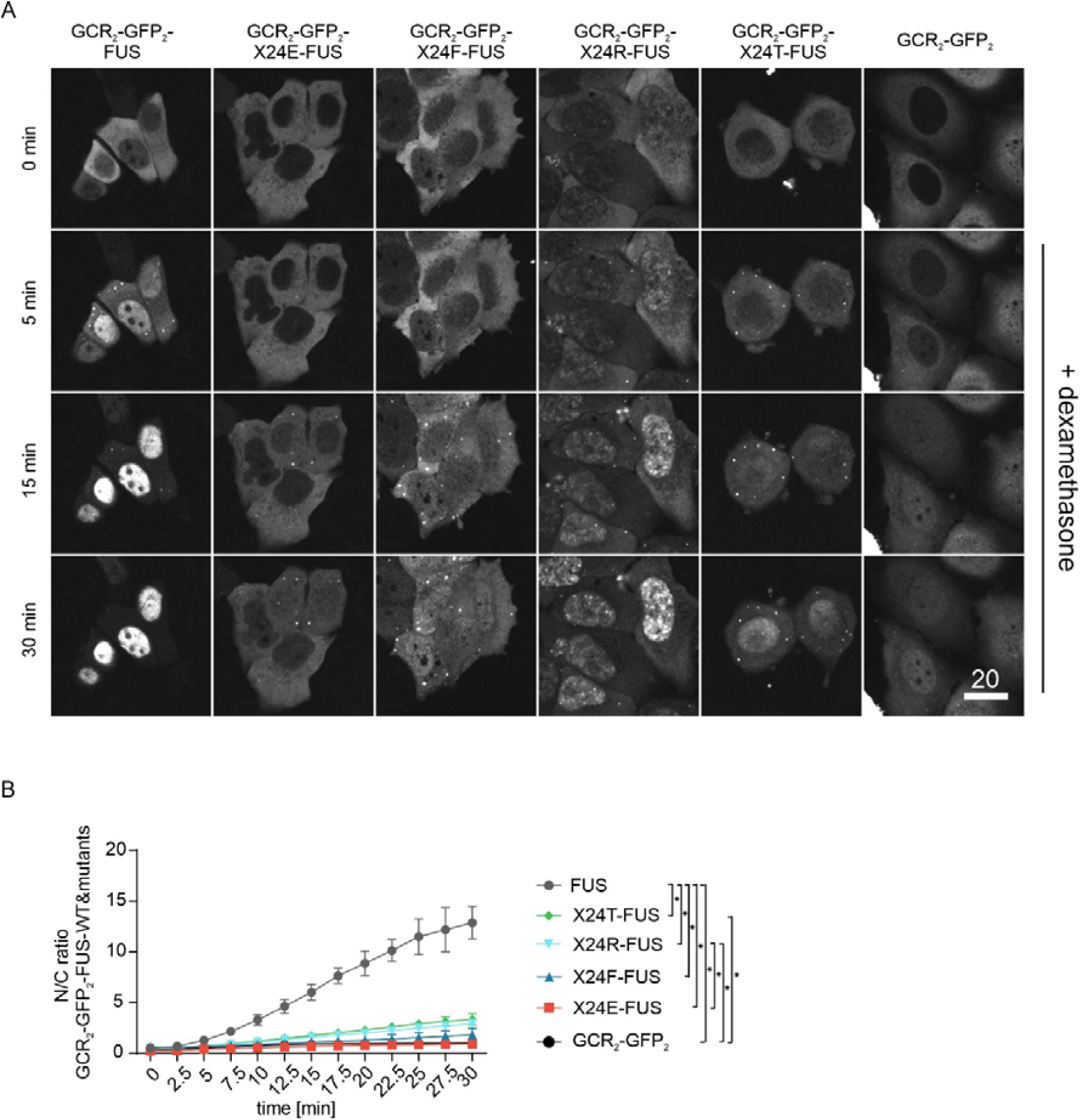
FUS mutants interfere with the nuclear localization of FUS in intact HeLa cells. (A) Hormone-inducible nuclear import assay in cells transfected with either GCR_2_-GFP_2_-FUS, GCR_2_-GFP_2_-X24T-FUS, GCR_2_-GFP_2_-X24R-FUS, GCR_2_-GFP_2_-X24F-FUS or GCR_2_-GFP_2_-X24E-FUS. Cells transfected with a GCR_2_-GFP_2_ reported served a diffusion control. Scale bar, 20 µm. (B) Nuclear import rates are plotted as nuclear/cytoplasmic (N/C) ratio of fluorescence intensities of all FUS reporters shown over time as the mean of three independent experiments ± SD. Statistical significance was calculated by repeated-measures two-way ANOVA with Tukey’s multiple comparison tests and significant differences between different groups only showed at 30 min time points (*P <0.05).

## DISCUSSION

Condensate formation plays an important role in many physiological processes, such as transcription, genome organization, immune response, and neuronal synaptic signaling. Dysregulation of condensate formation can disrupt nucleocytoplasmic trafficking, cause protein aggregation, and disrupt cellular homeostasis, leading to neurodegenerative disease and cancer (64,65). The formation of condensates often involves disordered RG/RGG regions, as well as other key players such as RNA and chaperones. The residues surrounding the RG/RGG regions are quite variable, and their role in phase separation, TNPO1 binding, and chaperoning is still not fully understood. In this study, we demonstrated how the sequence context within RG/RGG regions regulates their propensity for phase separation, TNPO1-binding *in vitro*, recruitment to SGs, and nuclear import of cargo proteins in cells.

Starting with a systematic approach and synthetic [XRGG]_7_ peptides, we combined coarse-grained slab simulations with in vitro turbidity and microscopy to disentangle how the residue immediately preceding RG tunes RNA-driven assembly. In peptide-only simulations, all [XRGG]_7_ variants remained largely mixed (K≈1), indicating that variation at the X position alone does not drive robust demixing under these conditions (Figure 1A). In contrast, addition of RNA uncovered strong sequence dependence: peptide partition coefficients spanned more than two orders of magnitude across the library, and RNA partitioning closely followed the same trend, consistent with co-condensation (Figure 1B). Representative systems illustrate the extremes, with [RRGG]_7_ forming a compact RNA-rich slab and [ERGG]_7_ remaining weakly partitioned without a distinct dense phase (Figure 1C–F). Guided by these results, turbidity assays showed that only a subset of [XRGG]_7_ sequences forms RNA-induced condensates in vitro, most prominently those with positively charged residues (Arg, Lys, His), the aromatic residue Tyr, and hydrophobic residues with long side chains (Leu, Met), with additional effects for selected polar residues (e.g., Cys/Asn/Ser) and Gly (Figure 2B–H; Supplementary Figure S1A–B). Together, these data support the view that RNA does not act as a generic driver, but rather amplifies intrinsic sequence biases at the X position into pronounced differences in condensate formation propensity. This view is supported by recent work on natural RG-rich domains showing RNA-triggered condensation of an arginine–glycine-rich coilin segment, together with broader evidence that RGG/RG-mediated RNA interactions rely on electrostatic and π-type contacts that can be strongly modulated by local sequence context (20,22,66). The [XRGG]_7_ residues can form several types of interactions, including electrostatic, cation-π, and hydrogen-bonding with multiple sites within RNA, including the negatively charged phosphate backbone, ribose sugar hydroxyls, and the aromatic rings (21,67,68). Interestingly, we found that, although that they show only a single-atom difference, [CRGG]_7_ was slightly more prone to form condensates than [SRGG]_7_. Although the precise molecular details of this difference remain to be elucidated, there are first indications that sulfur-containing amino acids, like cysteine, might engage in interactions with side chains of aromatic residues, as found in RNA, to form sulfur-π interactions (69,70). Additionally, Met and Cys sulfur moieties can form hydrogen bonds with NH groups of Arg guanidine, potentially contributing to the condensate formation (63–66). Except for [GRGG]_7_, [XRGG]_7_ peptides prone to phase separate have amino acids with side chains favoring the formation of multivalent interactions. Beyond the side chain properties, we found that the backbone flexibility plays an important role in condensate formation. Gly residues in RG/RGG regions increase flexibility, maintaining condensate formation (57,73), while Pro cyclic side chains reduce backbone flexibility and disturb secondary structure formation (74–76) In the [XRGG]_7_ background, the introduction of Pro residues abrogated condensate formation (Figure 2B–H). This suggests that condensate formation is highly sensitive to backbone flexibility as the remaining Gly residues were not sufficient to enable condensate formation in the RGG background. Moreover, we found that enriching negatively charged residues, such as aspartate or glutamate in the RGG background has a dominant suppressive impact on condensate formation (Figure 2B–H). This is in line with previous studies, for example by Bremer *et al.,* who showed that adding 12 aspartate or glutamate residues to the hnRNPA1-prion-like low-complexity domain weakens the driving forces for phase separation (77).

We analysed the human IDR dataset (Supplementary file 1) to place these minimal [XRGG]_7_ observations into a proteome-scale context and to define which RG/RGG organizations are most relevant *in vivo*. Our tract-level ‘RG-tract grammar’ is consistent with recent proteome-wide analyses showing non-random residue patterning around RG motifs and with sequence-based grammar models of RNA granules that emphasize short local features beyond motif counts as determinants of granule association (19,20). In line with our design choice, short RG/RGG repeat arrays predominate in human IDRs, with the majority of RG- and RGG-containing proteins carrying seven or fewer repeats in the searched spacing regime (Figure 2A). Beyond repeat number, our analysis further reveals an “RG-tract grammar”: in canonical RG/RGG proteins, RG motifs are typically concentrated into compact local tracts with short inter-RG linkers (most commonly ≤7 residues), whereas predicted disordered regions in general show a broader distribution of linker lengths and far fewer windows combining high RG density with very short spacers (Figure 3A–D). This architecture provides a straightforward route to multivalency, many closely spaced RG motifs presented in a small sequence window, and offers a mechanistic rationale for why even a single-residue change at the X position can have outsized effects once RNA is present: in compact tracts, small shifts in sticker chemistry are repeatedly “sampled” across many neighbouring motifs, effectively scaling up their impact on condensation. We analyzed the frequency of each amino acid within the human IDRs and RG-rich regions (Supplementary Figure S6A). Our analysis showed that evolution from IDRs to RG-rich regions retained aromatic residue, which are prone to either phase separate, and disfavored specific residues such as Asp (Supplementary Figure S6A). This suggests that evolution maintains a fine balance of RG/RGG regions of condensate-promoting aromatic residues and residues preventing condensate formation such as negatively charged aspartate. This is for example supported by the observation that mutations of aspartate residues in the RG/RGG regions of TAF15 PLD favor the phase separation (57). This suggests that the presence of Asp in RG-rich regions might be beneficial to prevent aggregate formation which is the later stage of phase separation. Furthermore, we performed separate analyses to identify the frequency of each residue within IDRs of proteins known to localize in SGs (Supplementary file 7) (39) (e.g., FUS, TAF15, EWSR1, hnRNPA1 and hnRNPA2) and within IDRs of proteins known to interact with TNPO1 (e.g., FUS, TAF15, hnRNPA1/2 and CIRBP) (40,41). We observed that retained residues within RG-rich regions were more pronounced within RG-rich regions in both the IDRs of proteins associated with SG localization (Supplementary file 7) and those binding to TNPO1 (Supplementary file 7) (Supplementary Figure S6A-C). In both subsets, hydrophobic residues (especially L/I/V/A) and Trp are depleted (log2 < 0) (Figure 3B,C), consistent with selection against sequence features that could favor irreversible self-association in vivo. In addition, we found that Asn and Ser, which exhibited lower tendencies for phase separation and had binding weaker affinities for TNPO1 binding, are retained in RG-rich regions of proteins known to localize to SGs and to bind TNPO1 (Supplementary Figure S6B and C). Ser residues are frequently found in PLCDs and function as spacers (77). Our observation hints at the potential role of Ser residues as spacers within RG-rich regions. Furthermore, Arg to Lys mutations disrupt phase separation of proteins (30,77). Mechanistically, this spacer/context interpretation aligns with recent evidence that neighbouring disordered sequence elements can reshape what core RGG/RG motifs do in RNA binding, and it matches proteome-wide context signatures indicating that residues flanking RG motifs are systematically biased rather than interchangeable (20,22). In line with those findings, we demonstrated that Lys residues are depleted from RG-rich regions (Supplementary Figure S6A-C). Moreover, we observed that negatively charged Glu residues occur less frequently within RG-rich regions compared to IDRs, whereas negatively charged Asp residues are retained in RG-rich regions (Supplementary Figure S6A-C). It has been shown that the exclusion of four Asp residues from A1-LCD decreased propensity for phase separation although the total net charge of protein increased. This suggested that Asp can enhance phase separation in specific sequence contexts, which aligns with our analysis showing the presence of Asp in RG-rich regions. Previous research has showed that introducing 12 Glu mutations in A1-LCD wild-type resulted in a slightly higher saturation concentration compared to 12 Asp mutations because of the higher steric bulk of Glu over Asp (77). This suggested that Glu has a stronger effect in preventing phase separation, which is consistent with our analysis showing a depletion of Glu residues in RG-rich regions (Supplementary Figure S6A-C).

In this work, we furthermore aimed to reveal how the sequence context within the RG/RGG regions affects TNPO1 binding and chaperoning. We found that [XRGG]_7_ peptides with the X position containing Arg, Lys, Tyr, His, Leu and Met bind to TNPO1 with nanomolar affinities, comparable to the affinities reported for the classical PY-NLS (28,78)^24, 68^. No TNPO1-binding was detected for the negatively charged peptides [DRGG]_7_ and [ERGG]_7_ by ITC. Our previous study showed that in addition to the classic PY-NLS binding site, RG/RGG regions can bind to TNPO1 via a disordered negatively charged loop and a negatively charged surface localized adjacent to the PY-NLS binding site (28,78–81). In the minimal [XRGG]_7_ context lacking a PY-NLS, enrichment of negatively charged residues (like glutamate and aspartate) is expected to weaken electrostatic complementarity and can reduce detectable TNPO1 binding in our assays (such as [ERGG]_7_, [DRGG]_7_). However, this trend does not directly extrapolate to FUS-RGG-PY constructs, where the presence of the PY-NLS module changes the binding context and X6E does not reduce TNPO1 affinity, indicating that TNPO1 recognition reflects combined contributions from multiple elements rather than a single “charge grammar.”

Inspired by the observation that TNPO1 can bind to RG/RGG regions of FUS and CIRBP, and suppress their phase separation (28,30–32), we studied TNPO1 chaperoning of [XRGG]_7_ condensate formation by turbidity assays and DIC microscopy (Figure 5A-E). Summarizing, we found that the sequence context has a strong impact on TNPO1-mediated chaperoning and that the chaperoning effect is strongly linked to TNPO1 binding affinity. This indicates that TNPO1-binding strength and TNPO1-mediated chaperoning of RG/RGG proteins co-evolved with of condensate-formation in order to provide cells with a regulatory mechanism preventing the adverse effects of condensate formation, as observed in neurodegenerative diseases (28,30–32). This model aligns with recent disease-focused frameworks in which nuclear import receptors, prominently TNPO1, act as ‘gatekeepers’ that couple efficient nuclear localization to suppression of aberrant phase transitions of prion-like RBPs, and with experimental evidence that regulatory inputs within RGG regions (including arginine methylation) can rewire interaction networks and shift condensate behavior toward, or away from pathological state (27,34).

We further extended our studies beyond the [XRGG]_7_ model peptides and included RG/RGG regions of human proteins. Interestingly, and in line with previous studies, we found that the propensity for RNA-mediated condensate formation strongly depends on the charge density. Peptides with high positive charge density underwent phase separation, while peptides which a low positive charge density did not (Figure 6A-B and Table 2). Condensate formation seems to reach a plateau at a charge density of around +0.3. Beyond this, presence of additional aromatic residues did not further enhance condensate formation, as indicated by the similar behavior of PAIRB^363-387^ (no aromatic residues) compared to NUCL^653-695^, FBRL^8-47^, and LS14A^401-431^ (Figure 6A and Table 2). Furthermore, we found that the sole presence of aromatic residues was insufficient to explain condensate formation, as HNRPU^700-730^ and HNRL1^637-659^ did not form condensates upon addition of RNA. Analysis of the positions of aromatic residues within our selected natural RG/RGG sequences showed that aromatic residues in “FG”, “FR” or “YR” positions contributed to phase separation, as observed in NUCL^653-695^, FBRL^8-47^ and LS14A^401-431^, but not in “FN” or “FQ” positions at the end of the sequence, as seen in HNRPU^700-730^ and HNRL1^637-659^. FG sequences are enriched in nucleoporins and are known to promote condensate formations (82–85). Our data suggest that FG repeats in RG/RGG regions also contribute to phase separation. Strikingly, and in line with the data obtained for our model peptides, we discovered that TNPO1 bound and chaperoned all peptides, dependent on the charge density, with HNRL1^637-659^ showing the lowest charge density (+0.13) and no binding (Figure 6C-E, Supplementary Figure S3A-F, and Table 1-2). Given that the TNPO1 binding affinities of these peptides are comparable to the affinities described for CIRBP and FUS (28), we propose that all of the binders could be yet undiscovered TNPO1 cargoes.

We further validated our model using the FUS^RGG3-PY^ region and introduced six mutations, including Arg, Phe, Tyr, Thr, Cys and Glu at the [X] positions preceding the RG/RGG motif (Figure 7A). We found that aromatic residue mutations (Tyr and Phe) increased phase separation, whereas Arg mutations did not have an additive effect on phase separation (Figure 7B). Aromatic residues (Phe and Tyr) have been proposed to act as “stickers” in prion-like low complexity domains, to form intra- and intermolecular contacts and in turn promote phase separation, whereas Arg has been proposed to act as a context-dependent auxiliary sticker (77,86). Our findings suggest that Phe and Tyr act as “stickers” for RG/RGG regions as well and promote phase separation. In line with our data obtained for other RG/RGG proteins, there seems to be a limit beyond which an increase in charge density or aromatic content does not translate into additional phase separation. As expected, the introduction of negatively charged residues disrupted the phase separation of FUS^RGG3-PY^ in a dominant manner (Figure 7B). This is in agreement with previous work showing that interactions between the RG/RGG region of FUS and RNA are mediated by electrostatic interactions (87). Glutamate mutations in FUS^RGG3-PY^ decreased the total net charge of the mutant compared to wild-type FUS^RGG3-PY^ (Table 3). This suggests that low net charge reduces electrostatic interactions between FUS and RNA. Although the model peptides [TRGG]_7_ and [CRGG]_7_ showed striking differences in phase separation, we did not observe any substantial impact of these mutations in the natural FUS^RGG3-PY^ background (Figure 7B). This could be due to the fact that introducing Thr or Cys did not substantially affect charge density and the number of stickers. Our results showed that the individual effects of Thr and Cys in [XRGG]_7_ peptides can be considered neutral and are not able to dominate phase separation of FUS^RGG3-PY^.

When carrying out TNPO1 binding studies of FUS^RGG3-PY^ mutant peptides, we made the unexpected observation that arginine mutations decreased binding affinity even though the native PY-NLS remained intact (Figure 7C, Supplementary Figure S4A-G, and S5, and Table 1). This suggests that the highly positively charged RG/RGG can compete with the PY-NLS. In line with this, we and others showed that arginine-rich poly-GR and poly-PR repeat peptides bind TNPO1 and impair nuclear import of TNPO1 cargoes in C9orf72-ALS/FTD patients (7,88,89). In semi-permeabilized cells, we observed that arginine mutations significantly increased the recruitment of X6R_FUS^RGG3-PY^ to SGs (Figure 8C-D) suggesting enhanced RNA binding might be underlying recruitment into pre-formed SGs. Given that glutamate mutations abolished the recruitment to SGs (Figure 8C-D), we conclude that the sequence context of the RG/RGG region in FUS^RGG3-PY^ has a substantial impact on phase separation, TNPO1 chaperoning *in vitro*, and recruitment to pre-formed SGs in semi-permeabilized cells. Furthermore, our results indicate that the sequence composition of RG/RGG regions has evolved to well-balance and fine-tune its functional properties.

Finally, we performed nuclear import assays to test the effect of mutations on the nuclear import of the RNA-binding protein FUS. In line with our *in vitro* TNPO1-binding and chaperoning experiments, we observed that wild-type FUS localized to the nucleus and did not associate with any puncta (Figure 9A). Surprisingly, Thr, Arg, Phe and Glu mutations interfere with FUS nuclear import even in the presence of an intact PY-NLS (Figure 9A-B). These findings not only highlight the importance of the RG/RGG regions for FUS nuclear import and sub-cellular localization but further support our model that the sequence context of RG/RGG regions evolved to support their functional properties.

In summary, we showed that the sequence context of RG/RGG regions is important for their biophysical properties and nuclear localization. Our systematic analysis highlighted the variability of residues in RG/RGG regions in the human proteome. We observed that enrichment of RG/RGG regions with positively charged and/or aromatic residues enhanced phase separation, TNPO1-binding, and SGs recruitment, whereas negatively charged residues have opposite role. We showed that mutations in RG/RGG regions interfere with TNPO1-binding and nuclear import in intact cells, despite the presence of an intact PY-NLS. These findings highlight the importance of RG/RGG regions over an intact PYNLS in regulating the nuclear localization of FUS. Our findings provide important insight into the fundamental mechanisms controlling SG recruitment and nuclear import of RBPs with RG/RGG regions, and will help to comprehend cytoplasmic mislocalization of specific RBPs in the context of diseases.

## AUTHOR CONTRIBUTIONS

Sinem Usluer: Methodology, Validation, Formal analysis, Investigation, Data curation, Writing-original draft preparation, Visualization. Yukti Khanna: Conceptualization, Methodology, Software, Formal analysis, Investigation, Data curation, Writing-original draft preparation, Visualization. Saskia Hutten: Methodology, Validation, Formal analysis, Investigation, Data curation, Writing-review and editing, Visualization. Đesika Kolarić: Methodology, Validation, Formal analysis, Investigation, Data curation, Writing-review and editing. Benjamin Bourgeois: Conceptualization, Methodology, Validation, Formal analysis, Investigation, Writing-review and editing, Supervision. Iva Pritišanac: Validation, Writing-review and editing, Supervision. Dorothee Dormann: Validation, Writing-review and editing, Supervision, Funding acquisition. Tobias Madl: Conceptualization, Validation, Resources, Writing-review and editing, Supervision, Project administration, Funding acquisition.

## ACKNOWLEDGEMENTS

We thank the Center for Medical Research, Medical University of Graz, Graz, Austria for laboratory access, and thank Hansjörg Habisch (Medical University of Graz, Austria) for support with statistical analysis. We acknowledge support by the core facilities Bioimaging of the Biomedical Center Munich and Microscopy Core Facility of the IMB Mainz.

## Data and code availability

All scripts used for analysis and figure generation are archived on Zenodo (Y. Khanna, rg-rich-regions, DOI: 10.5281/zenodo.18375256) and are mirrored on GitHub (https://github.com/yukti-khanna/rg-rich-regions). Simulation data underlying the figures will be deposited on Zenodo and linked to the software record upon publication.

