## Supplementary Figures for "The sequence context of RG/RGG motifs determines condensate formation, transportin-1 binding and chaperoning"

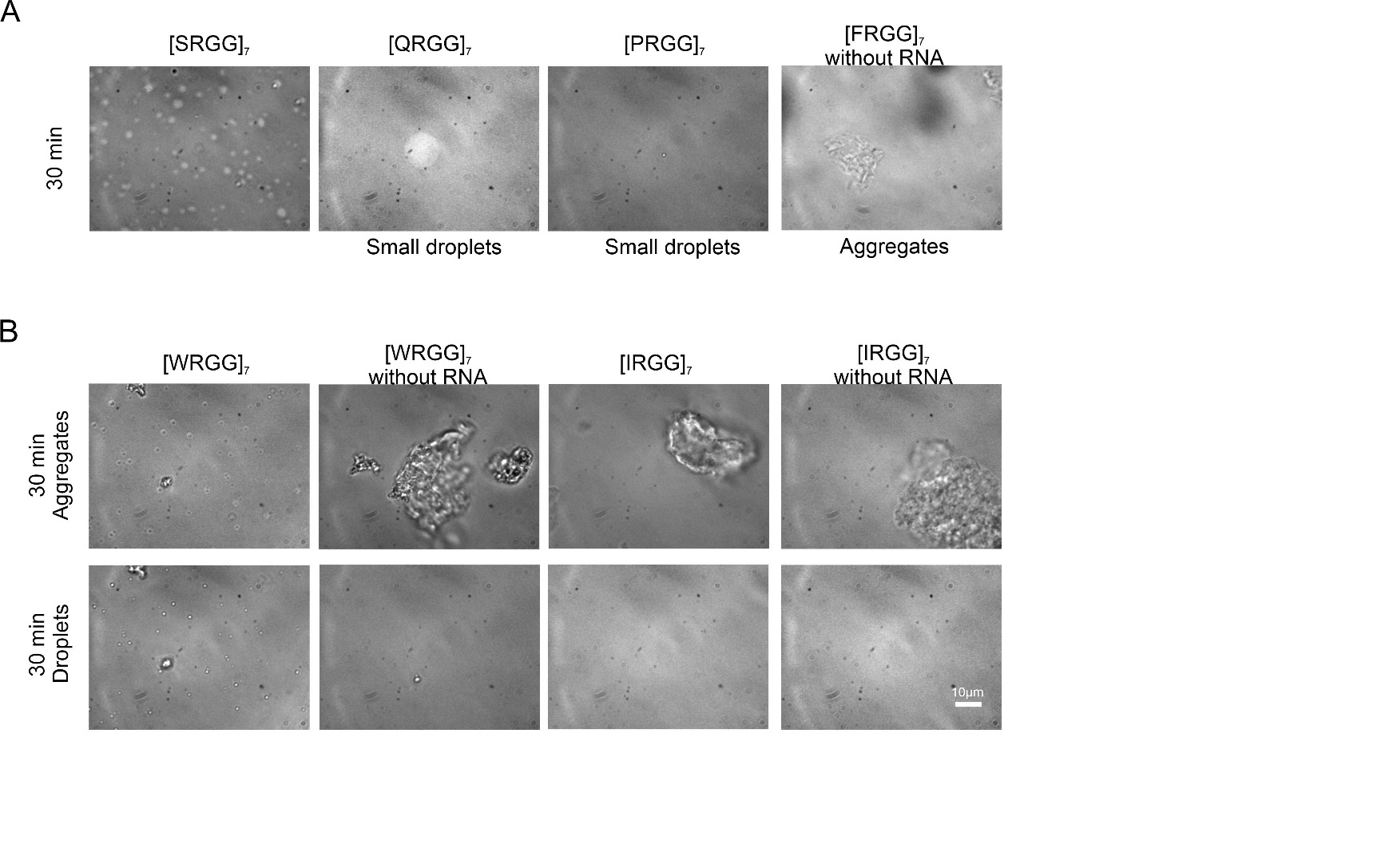

**Supplementary Figure S1:** Phase separation or aggregation of [XRGG]_7_ peptides. (A) DIC microscope images of [SRGG]_7,_ [QRGG]_7_, and [PRGG]_7_ peptides which form droplets and very small droplets, using peptides at 30 μM in presence of 15 μM RNA. In the right panel, DIC microscope images showing aggregates for [FRGG]_7_ in absence of RNA. (B) DIC microscope images of [WRGG]_7_ and [IRGG]_7_ peptides which form small droplets and aggregates, using peptide at 30 μM in the absence or presence of 15 μM RNA. Images were recorded 30 min after addition of RNA. (Scale bar: 10 μm)

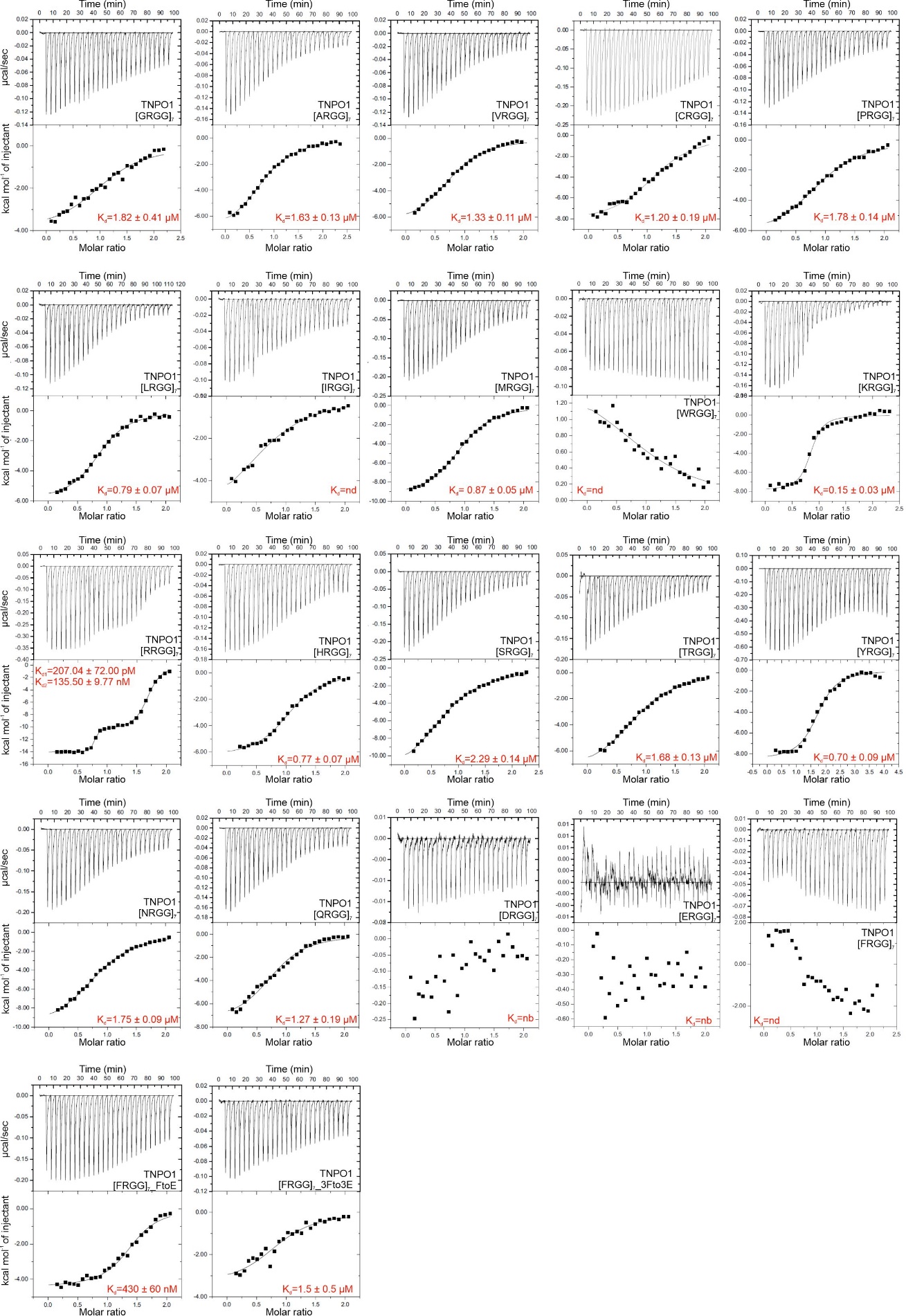

**Supplementary Figure S2:** ITC binding curves for [XRGG]_7_ peptides. Titration of 100 μM [XRGG]_7_ into 10 μM of TNPO1 except [YRGG]_7_. 200 μM [YRGG]_7_ was titrated into 10 μM of TNPO1. K_D_ values were indicated in the figure. The spacing time between each injection is 210 seconds, except [LRGG]_7_ which is 240 seconds. The reported errors correspond to the SD of the fit.

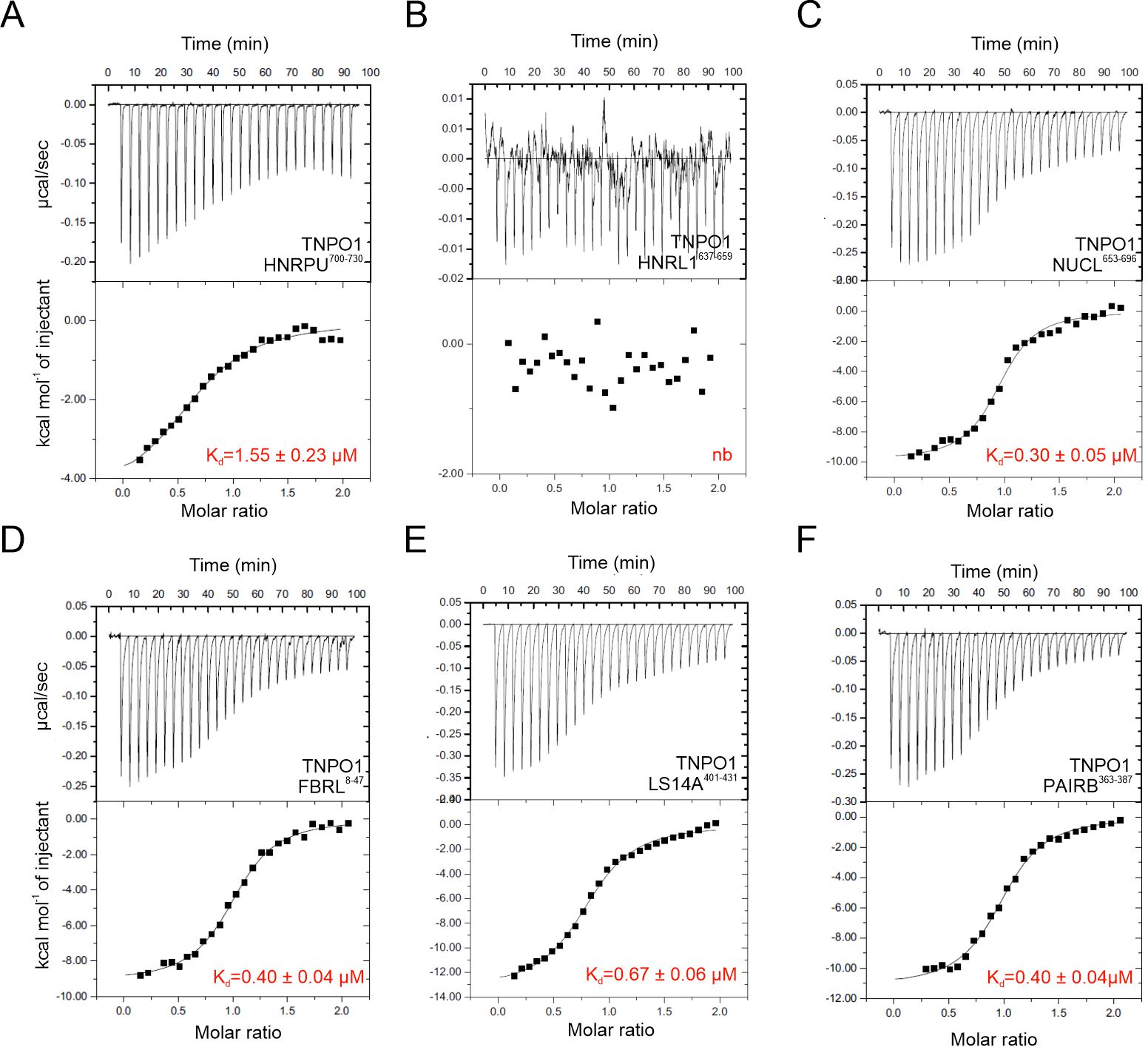

**Supplementary Figure S3:** ITC binding curves for RG/RGG-rich human peptides. (A-F) Titration of 100 μM RG/RGG-rich human peptides into 10 μM of TNPO1. K_D_ values were indicated in the figure. The reported errors correspond to the SD of the fit.

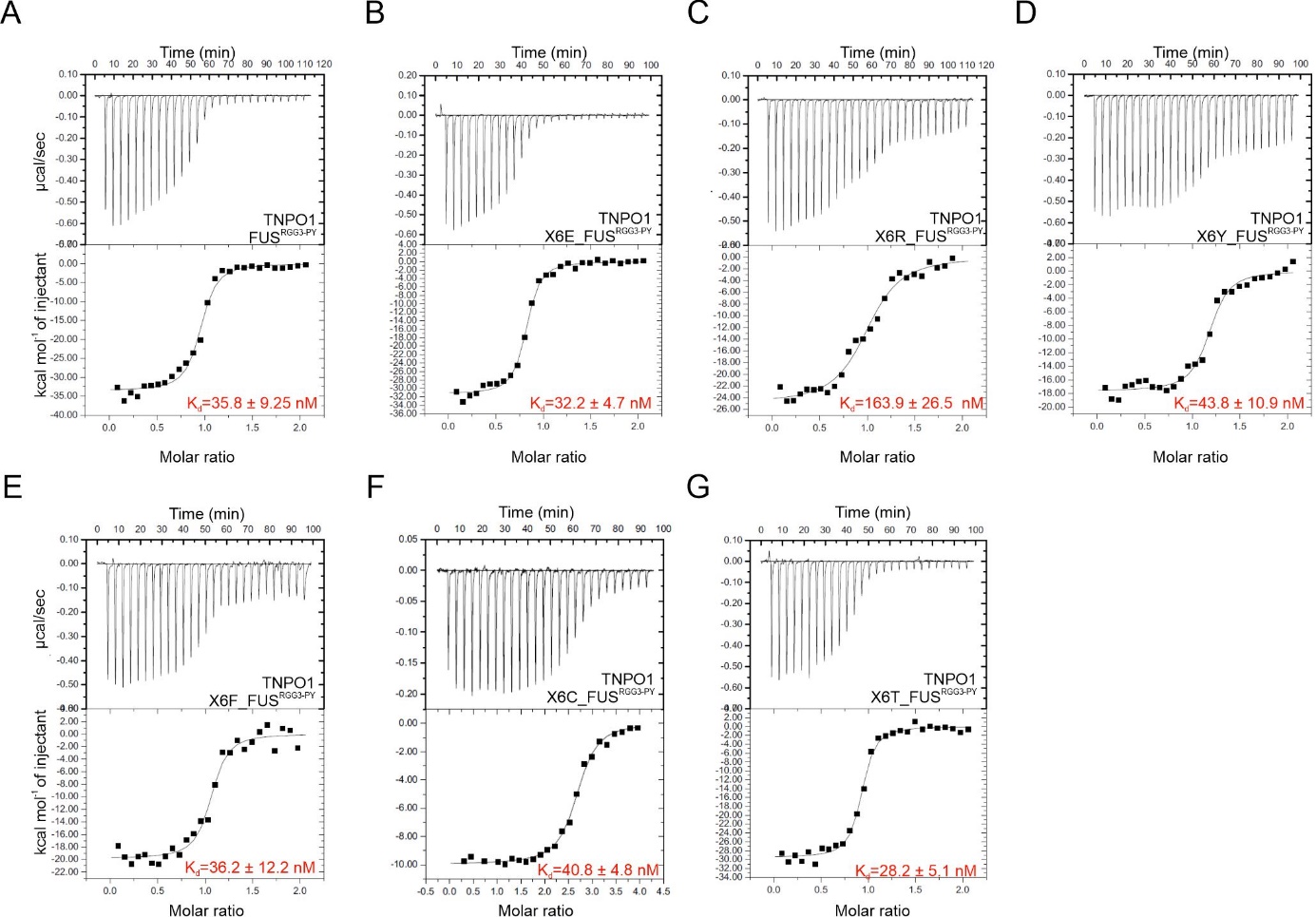

**Supplementary Figure S4:** (A-G) Titration of 50 μM FUS^RGG3-PY^ and mutants into 5 μM of TNPO1, except X6C_FUS^RGG3-PY^. 50 µM C mutant was titrated into 2.5 µM TNPO1. K_D_ values were indicated in the figure. The spacing time between each injection is 210 seconds, except for FUS^RGG3-PY^ and X6R_ FUS^RGG3-PY^ which are 240 seconds The reported errors correspond to the SD of the fit.

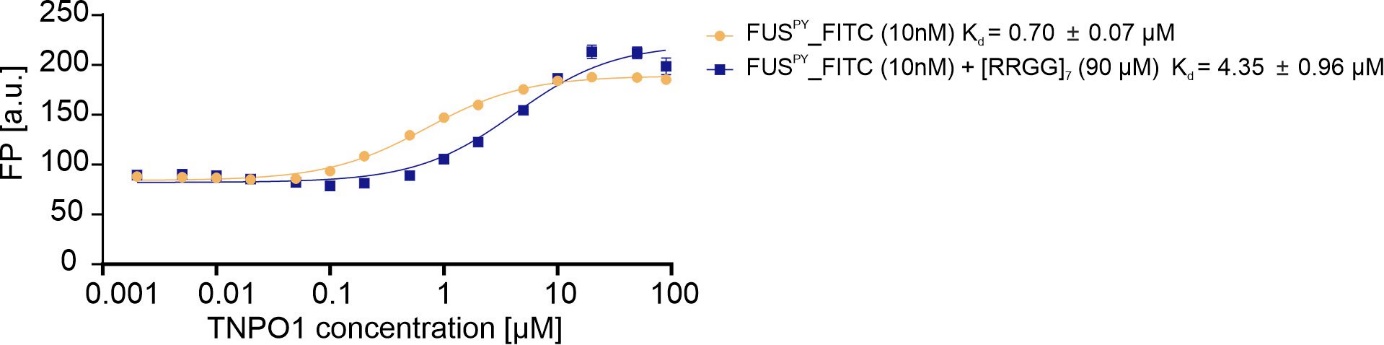

**Supplementary Figure S5:** [RRGG]_7_ peptide interferes with the binding of FUS^PY^ to TNPO1. Fluorescence polarization (FP) measurements of fluorescein isothiocyanate (FITC)-labeled FUS^PY^ (bisque) or FITC-labelled FUS^PY^ with 90 µM [RRGG]_7_ (navy) with increasing concentrations of TNPO1. K_D_ values are shown and were calculated by assuming a 1:1 complex formation. The graph represents the mean of three experiments and the reported errors correspond to the SD of the fit.

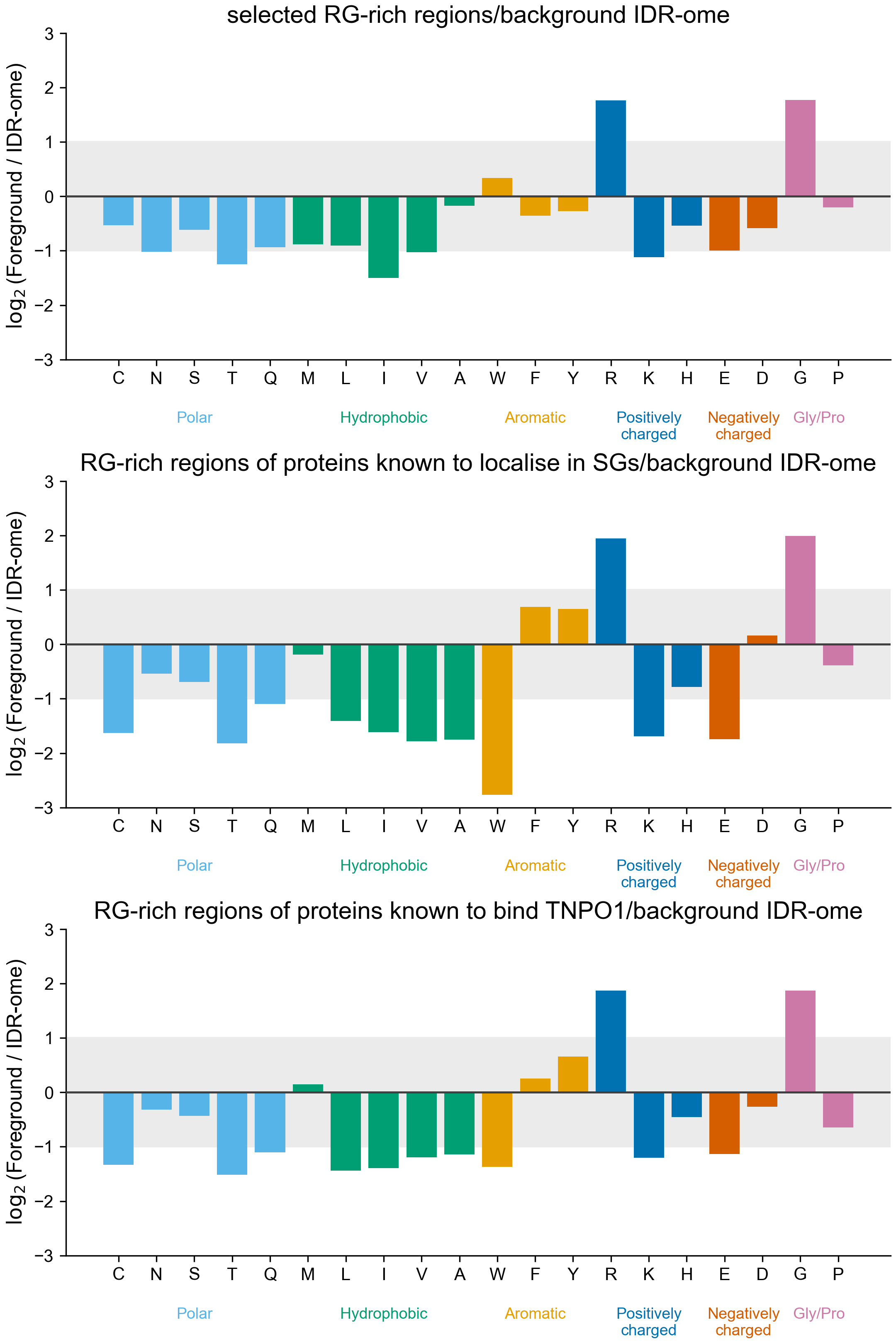

A

B

C

**Supplementary Figure S6:** Amino-acid enrichment profiles of RG-rich regions and their overlaps with PY and TNPO1 cargo proteins.

Bar plots show the log2 enrichment of each amino acid in the indicated foreground set relative to the background IDR-ome (Supplementary file 1). Foreground sets are: (B) selected RG-rich regions (Supplementary file 6), (2) RG-rich regions belonging to proteins in the PY list (Supplementary file 6, 7), and (3) RG-rich regions belonging to proteins in the TNPO1 cargo list (Supplementary file 6, 7). Amino acids are ordered by physicochemical class (polar, hydrophobic, aromatic, positively charged, negatively charged, glycine/proline), with group labels coloured to match the corresponding bars. The shaded band indicates the interval −1 to +1 (log2 units), highlighting modest deviations from background composition. Y-axis limits are fixed (−3 to +3) to enable direct comparison across panels.
